# A Single Cell Transcriptomic Atlas Characterizes Aging Tissues in the Mouse

**DOI:** 10.1101/661728

**Authors:** The Tabula Muris Consortium, Angela Oliveira Pisco, Aaron McGeever, Nicholas Schaum, Jim Karkanias, Norma F Neff, Spyros Darmanis, Tony Wyss-Coray, Stephen R Quake

**Affiliations:** Chan Zuckerberg Biohub, San Francisco, California, USA; Stanford University School of Medicine, Stanford, California, USA

## Abstract

Aging is characterized by a progressive loss of physiological integrity, leading to impaired function and increased vulnerability to death^1^. Despite rapid advances over recent years, many of the molecular and cellular processes which underlie progressive loss of healthy physiology are poorly understood^2^. To gain a better insight into these processes we have created a single cell transcriptomic atlas across the life span of Mus musculus which includes data from 23 tissues and organs. We discovered cell-specific changes occurring across multiple cell types and organs, as well as age related changes in the cellular composition of different organs. Using single-cell transcriptomic data we were able to assess cell type specific manifestations of different hallmarks of aging, such as senescence^3^, genomic instability^4^ and changes in the organism’s immune system^2^. This Tabula Muris Senis provides a wealth of new molecular information about how the most significant hallmarks of aging are reflected in a broad range of tissues and cell types.

---

We performed single cell RNA sequencing on more than 350,000 cells from male and female C57BL/6JN mice belonging to six age groups ranging from one month (human early childhood equivalent) to thirty months (human centenarian equivalent) (Figure 1a). We prepared single cell suspensions of the bladder, bone marrow, brain (cerebellum, cortex, hippocampus and striatum), fat (brown, gonadal, mesenteric and subcutaneous), heart and aorta, kidney, large intestine, limb muscle and diaphragm, liver, lung, mammary gland, pancreas, skin, spleen, thymus, tongue and trachea for all mice. Data were collected for all six age groups using microfluidic droplets (droplet), while the 3m, 18m and 24m time points were also analyzed using single cells sorted in microtiter well plates (FACS) (Extended Data Figure 1; Supplementary Tables 1&2; Supplementary Figures 1-3). Due to technical constraints, not every tissue was analyzed across all timepoints; for a complete list refer to Extended Data Figure 2a. The droplet data allow large numbers of cells to be analyzed using 3’ end counting, while the FACS data allow for higher sensitivity measurements over smaller numbers of cells as well as sequence information across the entire transcript length. Analyzing multiple organs from the same animal enables data controlled for age, environment, and epigenetic effects.

**Figure 1.**
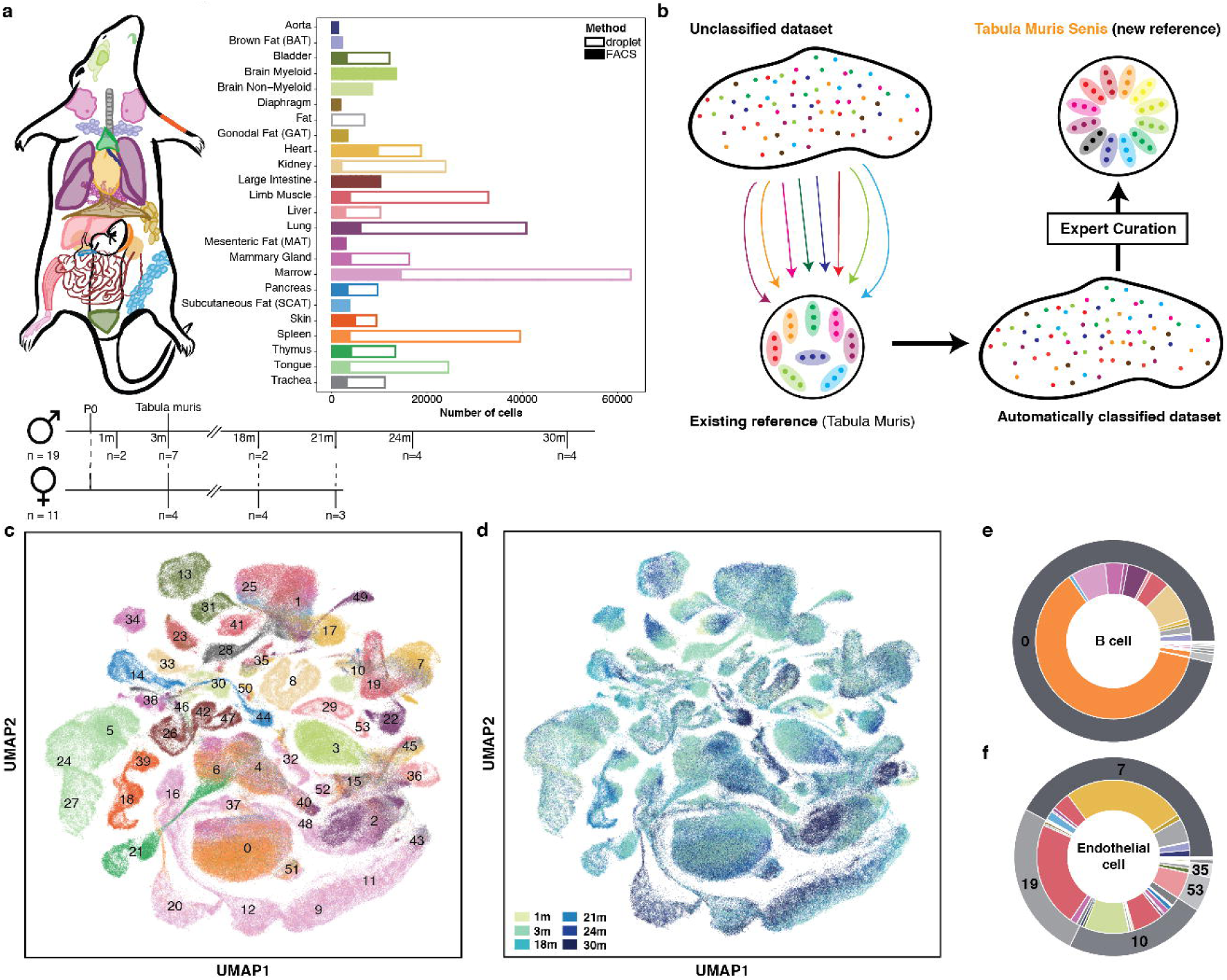
Overview of Tabula Muris Senis. **a**, 23 organs from 19 male and 11 female mice were analyzed at 6 different time points. The bar plot shows the number of sequenced cells per organ prepared by FACS (n=23 organs) and microfluidic droplets (n=16 organs). For the droplet dataset the Fat subtissues were processed together (Fat = BAT+GAT+MAT+SCAT). **b**, Annotation workflow. Data were clustered together across all time points. We used the Tabula Muris (3m time point) as a reference for the automated pipeline and the annotations were manually curated by tissue experts. **c,d**, UMAP plot of all cells, colored by organ and overlaid with the Louvain cluster numbers (**c**) and age (**d**); n = 356,213 individual cells. For the color dictionaries please refer to Extended Data Figure 2c. **e**, B cells (top) and endothelial cells (bottom) independently annotated for each organ cluster together by unbiased whole-transcriptome Louvain clustering, irrespectively of the organ they were found.

The previously published 3m time point, referred to as the *Tabula Muris*^5^, represents ∼20% of the cells in the entire dataset and was used as a basis to perform semi-automated cell type annotation of the additional time points (Figure 1b, Extended Data Figure 2b). Using this approach, we were able to automatically annotate over 70% of the cells. All the automated cell annotations were reviewed and approved by human experts, and the remaining cells were annotated by hand, creating one of the largest manually curated single cell transcriptomic resources in existence. Many of these cell types have not previously been obtained in pure populations, and these data provide a wealth of new information on their characteristic gene-expression profiles. Out of 529,823 total cells sequenced, 110,824 cells for FACS and 245,389 cells for droplet passed our strict filtering criteria (Extended Data Figure 2b) and were subsequently annotated (Extended Data Figure 1c,f), separately for each tissue and method; the remaining cells are also included in the on-line data set but were not used for further analysis here. To investigate whether cell annotations were consistent across the entire organism, we used bbknn^6^ to correct for method-associated batch effects. Following batch correction, we clustered all cells using an unbiased, graph-based clustering approach^7, 8^ (Figure 1c,d) and assessed the co-occurrence of similarly annotated cells in the same clusters. For example, cells annotated as B cells or endothelial cells tend to occupy the same clusters irrespectively of their tissue of origin or method with which they were processed (Figure 1e,f; Extended Data Figure 3).

Tabula Muris Senis provides a powerful resource with which to explore aging related changes in specific cell types. The entire dataset can be explored interactively at <u>tabula-muris-senis.ds.czbiohub.org</u>. Gene counts and metadata are available from figshare (https://figshare.com/projects/Tabula_Muris_Senis/64982) and GEO (GSE132042), the code used for the analysis is available from GitHub (https://github.com/czbiohub/tabula-muris-senis) and the raw data are available from a public AWS S3 bucket (https://s3.console.aws.amazon.com/s3/buckets/czb-tabula-muris-senis/). An important use of the single cell data is to resolve whether gene expression changes observed in bulk experiments are due to changes in gene expression in each cell of the population, or whether the gene expression in each cell stays constant but the number of cells of that type changes, or both. In a global analysis of gene expression changes using the Tabula Muris Senis and bulk RNAseq from tissues^9^, we observed that in many cases changes in gene expression are due to both changes in the numbers of cells in a population and to changes in the gene expression levels in each cell (Extended Data Figure 4). As a specific example of this approach, we investigated how the expression of Cdkn2a changes with age. As Cdkn2a/p16 is one of the most commonly used markers of senescence^10^ and an important hallmark of aging^11^, we computed the fraction of cells expressing Cdkn2a at each age. The fraction of cells expressing the gene more than doubled in older animals in both FACS (Figure 2a) and droplet (Figure 2b), accompanied by a 2-fold increase in the actual expression level of p16 by those cells that did express it (Figure 2c,d). It is worth noting that the fraction of cells expressing p16 in the 30m mice is smaller than at 24m, prompting us to speculate that perhaps the animals that live longest somehow have a slower rate of senescence. We next compiled a list of previously characterized senescence markers^12–15^ and plotted the fraction of cells expressing each marker across all age groups (Supplementary Table 3). Out of these markers, Cdkn2a has the highest correlation between aging and fraction of cells expressing the gene. Other genes for which the fraction of cells expressing significantly increased with age include E2f2^16^, Lmnb1^17, 18^ and Tnf and Itgax^19^. For some genes the fraction of cells expressing decreased with age, including members of the Sirt family (Sirt3, Sirt4 and Sirt5); this is consistent with previous literature finding that sirtuin is essential in delaying cellular senescence^20, 21^.

**Figure 2.**
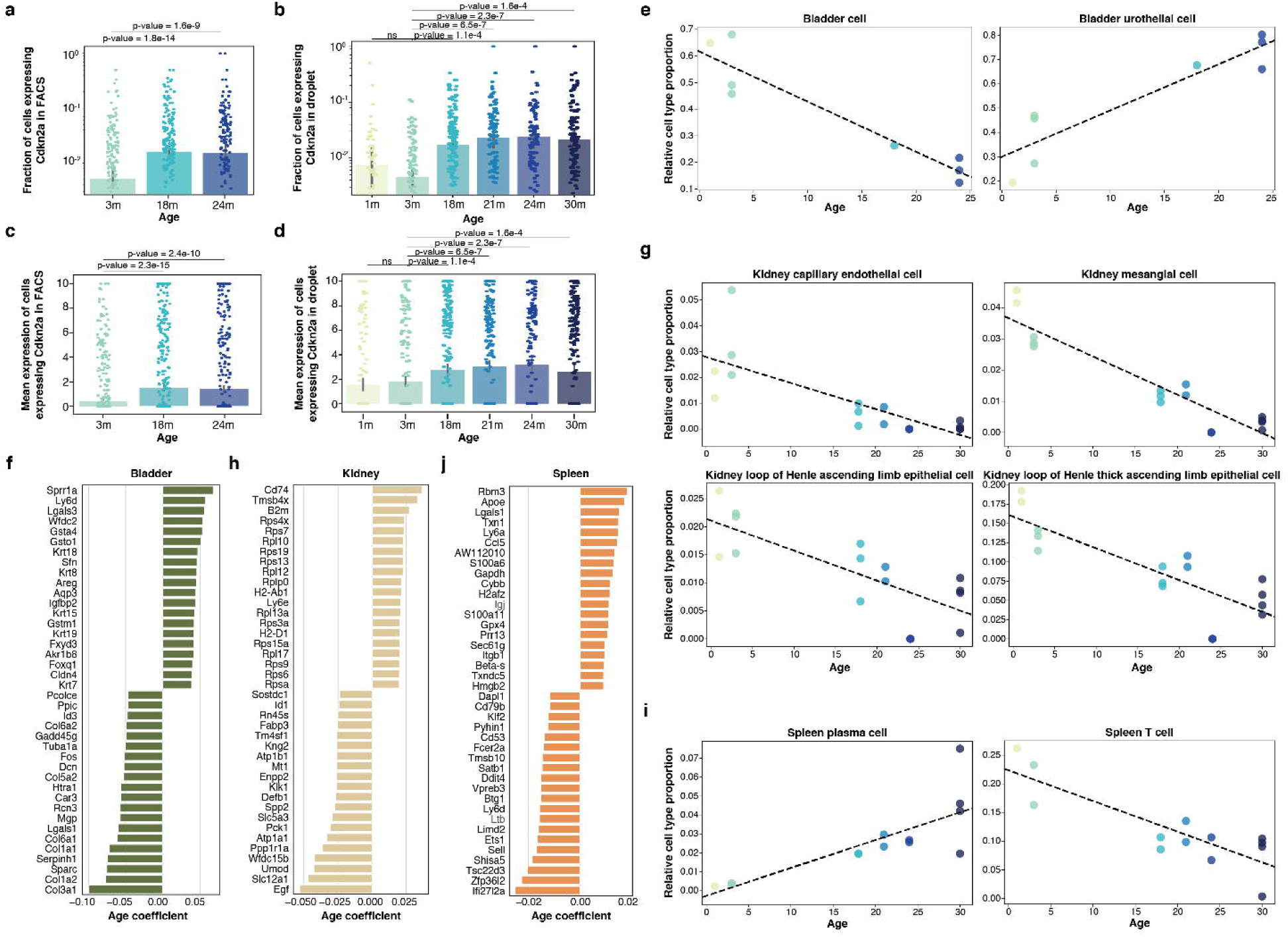
Cellular changes during aging. **a,b**, Bar plot showing the fractions of cells expressing Cdkn2a at each age group for FACS (**a**) and droplet (**b**). **c,d**, Bar plot of the median expression of Cdkn2a for the cells that do express the gene at each age group for FACS (**c**) and droplet (**d**). The p-value was obtained using a Mann-Whitney-Wilcoxon rank-sum two-sided test. **e**, Bladder cell (**left**) and bladder urothelial cell (**right**) relative abundances change significantly with age (p-value<0.05 and r^2^>0.7 for a hypothesis test whose null hypothesis is that the slope is zero, using two-sided Wald Test with t-distribution of the test statistic). **f**, Top 20 upregulated and downregulated genes in bladder using age as a continuous covariate while controlling for sex and technology. Genes were classified as significant under an FDR threshold of 0.01 and an age coefficient threshold of 0.005 (corresponding to ∼10% fold change). **g**, Kidney capillary endothelial cell (**top-left**), mesangial cell (**top-right**), loop of Henle ascending limb epithelial cell (**bottom-left**) and loop of Henle thick ascending limb epithelial cell (**bottom-right**) relative abundances change significantly with age (p-value<0.05 and r^2^>0.7 for a hypothesis test whose null hypothesis is that the slope is zero, using two-sided Wald Test with t-distribution of the test statistic). **h,** Top 20 upregulated and downregulated genes in kidney using age as a continuous covariate while controlling for sex and technology. Genes were classified as significant under an FDR threshold of 0.01 and an age coefficient threshold of 0.005 (corresponding to ∼10% fold change). **i**, Spleen plasma cell (**left**) and T cell (**right**) relative abundances change significantly with age (p-value<0.05 and r^2^>0.7 for a hypothesis test whose null hypothesis is that the slope is zero, using two-sided Wald Test with t-distribution of the test statistic). **j,** Top 10 upregulated and downregulated genes in spleen using age as a continuous covariate while controlling for sex and technology. Genes were classified as significant under an FDR threshold of 0.01 and an age coefficient threshold of 0.005 (corresponding to ∼10% fold change).

We investigated how the cellular composition of each tissue changes with age by evaluating how the relative cell type proportions within a tissue change with age. The overall cell composition for all tissues is in Extended Data Figure 5. While we present the cell composition for all tissues for which we have droplet data available, we only investigated the changes with age for tissues which had more than two time points (Supplementary Table 5). When interpreting compositional data, one must bear in mind that dissociation does not affect all of the cell types in a tissue equally, so changes in the relative composition of a given cell type with age are more meaningful than trying to compare proportions of different cell types at a single age^22–24^. Nonetheless, the changes in relative proportion of cell types provide important information on the effects of aging in a variety of tissues.

The bladder has pronounced changes in cell type composition with age (Figure 2e). While the mesenchymal compartment of this tissue decreases by a factor of three over the lifetime of the mouse (Figure 2e left), the urothelial compartment increases by a similar amount (Figure 2e right). The observation that the bladder urothelial cells increase with age is concordant with known age-related urothelial changes^25^. Differential gene expression analysis of overall tissue changes with age revealed that stromal-associated genes (Col1a1, Col1a2, Col3a1, Dcn) are downregulated while epithelial-associated genes (Krt15, Krt18, Sfn) are upregulated, supporting the compositional observations (Figure 2f; Supplementary Table 6). The decline of the endothelial population suggests that bladder aging in mice may be associated with lower organ vascularization, consistent with recent findings^26, 27^ and with the observed downregulation of vasculature associated genes Htra1 and Fos (Figure 2f; Supplementary Table 6). The increase in the leukocyte population could be indicative of an inflammatory tissue microenvironment, a common hallmark of aging which is consistent with literature on overactive bladders^28^ and supported by a significant overexpression of Lgals3, Igfbp2 and Ly6d across the tissue (Figure 2f; Supplementary Table 6) and by the overexpression of immune response associate genes such as Tnfrsf12a and Cdkn1a, by both bladder (mesenchymal) cells and bladder urothelial cells (Supplementary Table 6). Moreover, when comparing across ages, we observed that old leukocytes show increased expression of pro-inflammatory markers, such as Cd14, Lgals3 and Tnfrsf12a, and decreased expression of anti-inflammatory ones, such as Cd9 and Cd81, corroborating our hypothesis (Supplementary Table 6).

Age-dependent changes in the kidney include a decrease in the relative abundance of mesangial cells, capillary endothelial cells, loop of Henle ascending limb epithelial cells and loop of Henle thick ascending limb epithelial cells (Figure 2g). Both mesangial cells and capillary endothelial cells are core glomerular cells and their relative abundances reduction (Figure 2g top panels), together with downregulation of Egf and Atp1a1 (Figure 2h; Supplementary Table 6) suggest impaired glomerular filtration rate^29, 30^. This finding is reinforced by the differential gene expression results indicating that uromodulin (Umod), the most abundant protein in urine^31^, is downregulated. Umod is produced by the epithelial cells that line the thick ascending limb, and therefore given the relative decrease in the proportion of epithelial cells in the ascending and thick ascending limb, our results suggest that normal kidney functions are impaired^32^ (Figure 2g bottom panels, Figure 2h; Supplementary Table 6).

The liver is yet another tissue for which we observed changed tissue compositions with age, namely that the relative amount of hepatocytes decreases with age (Extended Data Figure 6a-d), which is supported by the reduction in the expression of albumin (Alb; Extended Data Figure 6e; Supplementary Table 6). Differential gene expression showed an increased immune signature, as illustrated by overexpression of H2-Aa, H2-Ab1, H2-D1, H2-Eb1, Cd74, Lyz2 and others (Extended Data Figure 6e). Previous findings suggested that pro-inflammatory macrophages drive cellular senescence and identified Il1b as a gene whose liver expression was remarkably different with age^12^ (Extended Data Figure 6f). We stained liver Kupffer cells (Extended Data Figure 6g) with Clec4f (canonical Kupffer cell marker) and found the number of Clec4f+ cells do not change with age, consistent with the results of the tissue composition analysis (Supplementary Table 7; Extended Data Figure 6h). However, when co-staining with Il1b, we found an increase with age in the number of cells expressing Clec4f and Il1b (Extended Data Figure 6h-j). Il1b has low expression in normal physiological conditions^33^. Specific blocking of IL1-RI (Il1b receptor) in hepatocytes has been shown to attenuate cell death upon injury, supporting the idea that increased expression of Il1b in Kupffer cells is typically a poor prognostic^34^. Regarding immune defense within the liver, sinusoidal endothelial cells (LSECs) play a unique role, being the main carriers of the mannose receptor (Mrc1) in the liver (Extended Data Figure 6k). Mrc1 expression in LSECs mediates endocytosis of pathogen and damage related molecules. Our findings identify increased Mrc1 age-related expression. Inflammatory signals have been found to up regulate Mrc1 expression and endocytosis ^35^. Staining for Mrc1 alongside classical LSEC marker Pecam1 (Supplementary Table 7; Extended Data Figure 6l) found the number of Mrc1 expressing LSECs increase over age (Extended Data Figure 6m-o). LSECs have a been found to have a reduced endocytic capacity in aged livers, while it has been suggested that LSECs proliferate after injury or that bone-marrow derived LSECs progenitors are recruited to the liver. This suggests that changes in LSEC gene signatures with age are linked closely with their function in immune response.

In the case of spleen our results show that with age the proportion of T cells decreases while the relative amount of plasma cells increases (Figure 2i). This is supported by upregulation of B cell/plasma cell markers (Cd79a, Igj; Figure 2j; Supplementary Table 6) and downregulation of Cd3d (Figure 2j; Supplementary Table 6). Similarly, in mammary gland we also observed a significant decline of the T cell population (Extended Data Figure 7a). Age-related decline of T cell populations has been associated with an increased risk of infectious disease and cancer^36^ and our results suggest that this may also happen in the spleen and mammary gland. We found that members of the AP1 transcription factors^37^ (Junb, Jund and Fos) were upregulated with age (Extended Data Figure 7b; Supplementary Table 6); this result is consistent with the observation that normal involution of the mammary gland is accompanied by significantly increased expression of many of these AP1-related transcription factors^38^.

Genomic instability is among the most widely studied aging hallmarks^1^ and the full-length transcript data from the FACS data allows the analysis of somatic mutation accumulation with age. We used the Genome Analysis ToolKit (GATK)^39^ to perform SNP discovery across all FACS samples simultaneously (Supplementary Table 8), using GATK Best Practices recommendations^40, 41^. We focused on genes expressed in at least 75% of cells for each age group within a particular tissue. We observed an age-related increase in the number of mutations across all of the organs we analyzed (Figure 3; Extended Data Figure 8a, 9a, 10a), with tongue and bladder being the most affected. Our analysis controls for sequencing coverage and gene expression levels (Extended Data Figure 8b, 9b, 10b). The number of mutations observed at each age are larger than technical errors due to amplification and sequencing errors, which can be estimated using ERCC controls that were spiked into each well of the microtiter plates^42^ (Figure 3; Extended Data Figure 8c-d, 9c-d, 10c-d). Despite the fact that it is difficult to infer genome-wide mutation rates from the transcriptome, which is known to inflate apparent mutational rates for a variety of reasons^42^, the observed trend is a useful indirect estimate of mutational frequency and genome stability.

**Figure 3.**
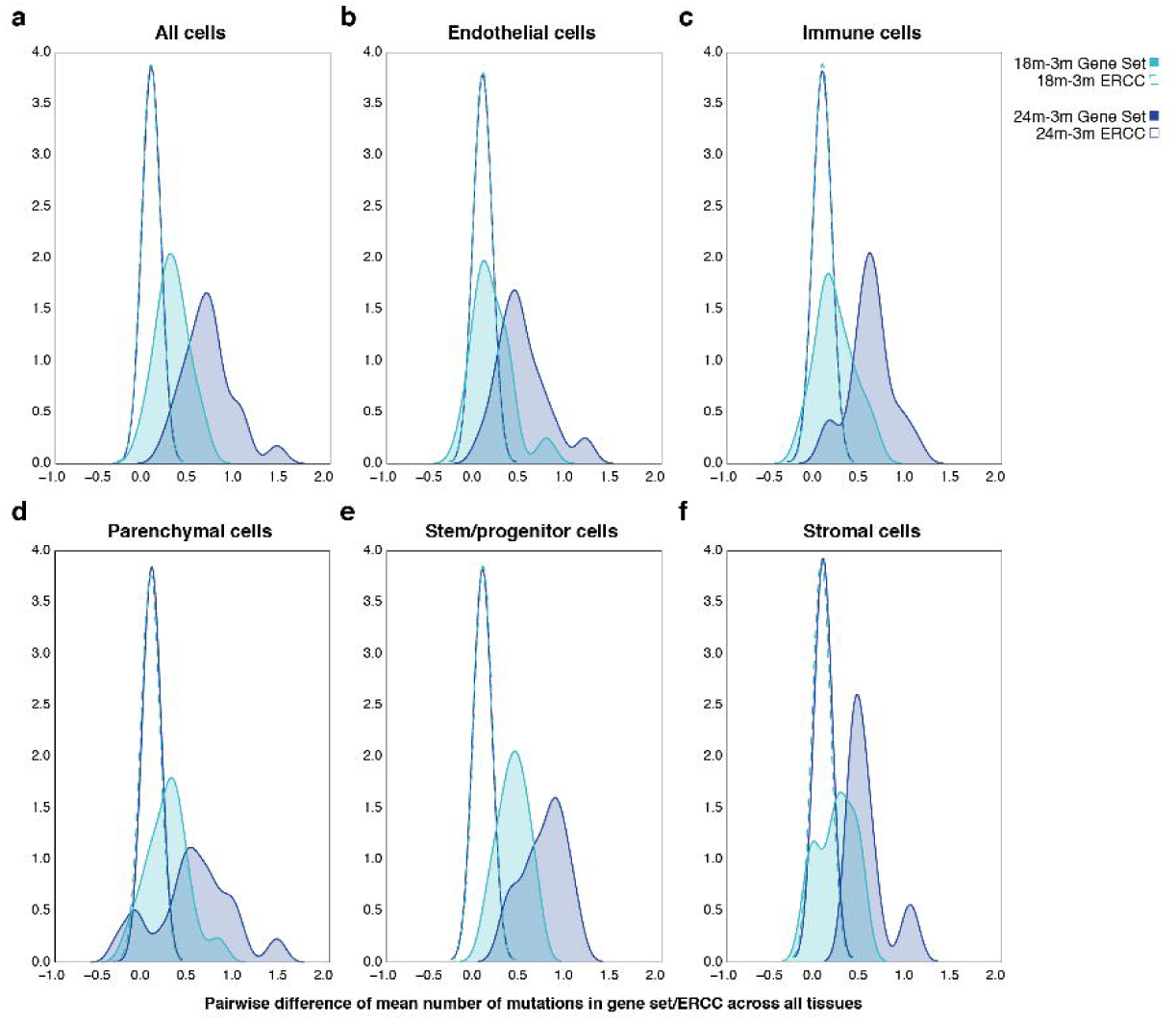
Mutational burden across tissues in the aging mice. Distribution of the difference of the mean mutation in the gene set (and ERCC spike-in controls) per cell between 24m and 3m and 18m and 3m for all tissues and cells (**a**) and with the cell types split in five functional groups, endothelial (**b**), immune (**c**), parenchymal (**d**), stem/progenitor cell (**e**) and stromal (**f**).

A final hallmark of aging which we investigated was the effect of age-induced changes on the immune system^2^. Analyzing a complete set of tissues from the same individual animal using the full-length transcripts obtained in the FACS data enabled us to analyze clonal relationships between B-cells and T-cells throughout the organism. We computationally reconstructed the sequence of the B-cell receptor (BCR) and T-cell receptor (TCR) for B cells and T cells present in the FACS data using singlecell-ige and TraCeR, respectively^43, 44^. BCRs were assembled for 6,050 cells (Figure 4a) and TCRs for 6,000 cells (Figure 4b). The number of cells with assembled BCRs was 1,818 for 3m, 1,356 for 18m and 2,876 for 24m old mice. We parsed the singlecell-ige^43^ output to define B-cell clonotypes based on the sequence of the assembled BCR (Supplementary Table 9) and found that while most of the cells at 3m were not part of a clone (9% were part of a clonal family), the number of B-cells belonging to a clonotype doubled at 18m (20%) when compared to 3m and doubled again from 18m to 24m (∼38%).

**Figure 4.**
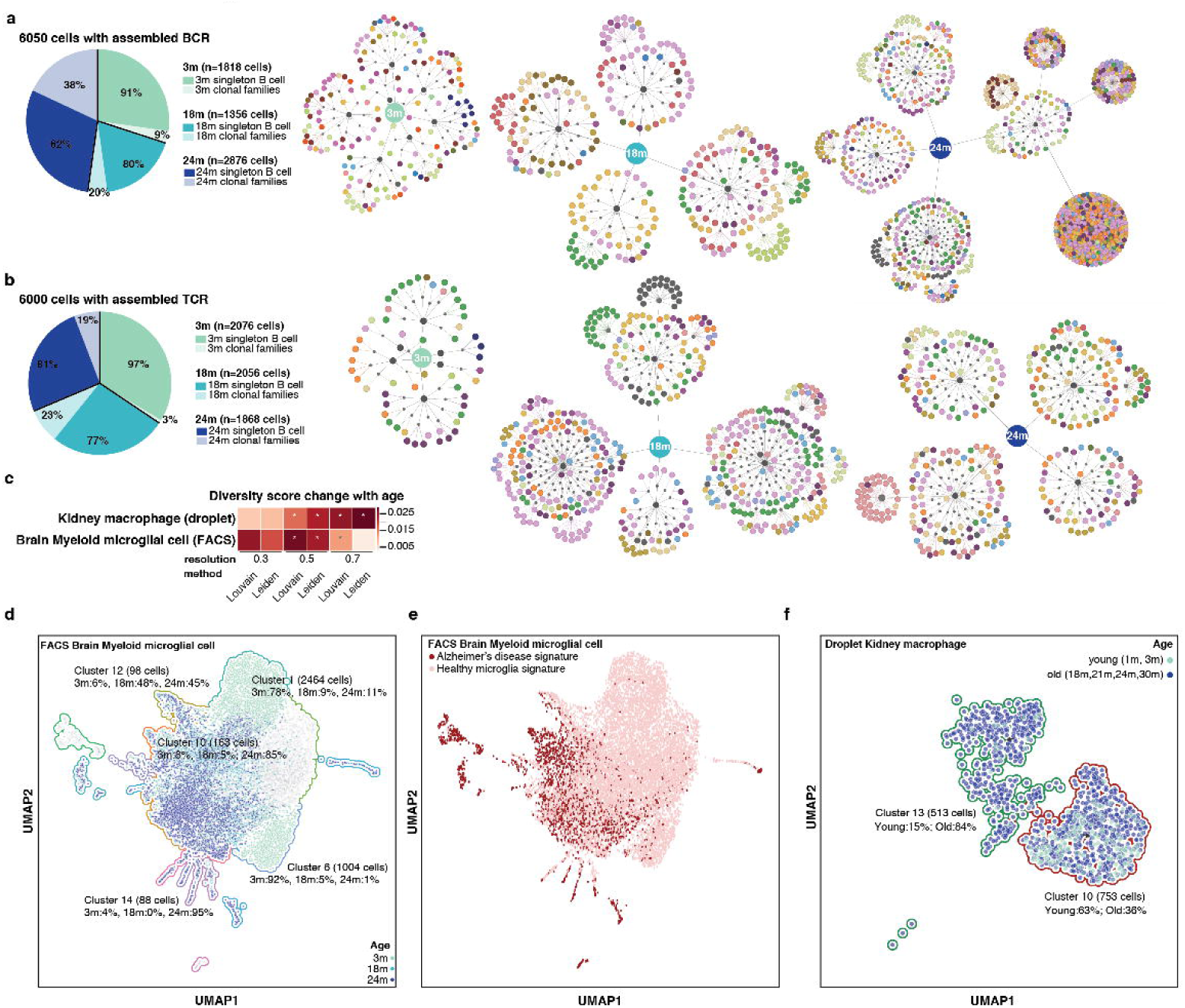
The aging immune system. **a**, B-cell clonal families. The pie chart shows the proportion of singleton B cells and B cells that are part of clonal families at 3m, 18m and 24m. For each time point, the clonal families are represented in a tree structure for which the central node is age. Connected to the age node there is an additional node (dark gray) that represents each animal and the clonal families are depicted for each animal. For each clonal family, cells that are part of that family are colored by the organ of origin. **b**, T-cell clonal families. The pie chart shows the proportion of singleton T cells and T cells that are part of clonal families at 3m, 18m and 24m. For each time point, clonal families are represented in a tree structure for which the central node is age. Connected to the age node there is an additional node (dark gray) that represents each animal and the clonal families are depicted for each animal. For each clonal family, cells that are part of that family are colored by the organ of origin. **c**, Diversity score for the two cell types that significantly change with age. **d**, UMAP plot of the brain myeloid microglial cell Leiden clusters (numbers) colored by age. Faded clusters do not change their relative age cell composition; colored clusters change their relative cell composition. **e**, UMAP plot of the brain myeloid microglial cells when scored using the microglia Alzheimer’s disease signature (Supplementary Table 10). **f**, UMAP plot of the kidney macrophage Leiden clusters (numbers) colored by age group.

The number of cells with assembled TCRs were roughly equal between 3m, 18m and 24m (2,076, 2,056 and 1,868 cells, respectively). Clonotype assignment is part of the output obtained by TraCeR^44^ (Supplementary Table 9). Interestingly, only ∼3% (55 out of 1,895) of the cells at 3m were part of a clone. For 18m, ∼23% (479 out of 2,056) of the cells were part of a clone and for 24m, ∼20% (348 out of 1,780) of the cells were part of a clone, indicating again an increase in clonality of the T-cell repertoire at later ages. These changes in clonality for both B and T cell repertoires are noteworthy because they suggest that the immune system of a 24m mouse will be less likely to respond to new pathogens, corroborating literature suggesting that older individuals have higher vulnerability to new infections and lower benefits from vacination^45, 46^.

As a final example of how the Tabula Muris Senis can be used to discover how cell types change with age, we computed an overall diversity score to identify which cell types were more susceptible to changes with age (Extended Data Figure 11). The diversity score is computed as the Shannon entropy of the cluster assignment and then regressed against age to provide a p-value (see Methods). We observed significant changes in diversity affecting cells of the immune system originating from the brain and in the kidney (Figure 4c, Extended Data Figure 12a,b). These results were not confounded by the number of genes expressed per cell (Extended Data Figure 12c,d). We found that in brain myeloid microglial cells, the majority of young (3m) microglia occupy clusters 1 and 6, while old (18m, 24m) microglia constitute the vast majority of cells in clusters 10, 12 and 14 (Figure 4d). Trajectory analysis suggests that young microglia go through an intermediate state, represented by the clusters mostly occupied by 18m microglial cells before acquiring the signature of old microglia (Extended Data Figure 12e). Clusters 10, 12 and 14 are mainly comprised of 18- and 24-month old microglia. These cells up-regulate MHC class I genes (H2-D1, H2-K1, B2m), along with genes associated with degenerative disease (e.g. Fth1)^47, 48^. When contrasting with clusters 1 and 6, which contain mostly 3m microglia, clusters 10, 12 and 14 gene expression is enriched with interferon responsive or regulatory genes (e.g. Oasl2, Oas1a, Ifit3, Rtp4, Bst2, Stat1, Irf7, Ifitm3, Usp18, Ifi204, Ifit2), suggesting an expansion of this small pro-inflammatory subset of microglia in the aging brain^49^. Moreover, the list of differentially expressed genes between “young” and “old” clusters resembled the Alzheimer’s disease specific microglial signature previously reported^47^, with 55 out of the top 200 differential expressed genes being shared between the two differential gene expression lists (Figure 4e; Supplementary Table 10). Regarding kidney macrophages, we found two clusters that remarkably changed their composition with age. Cluster 10 is primarily composed of cells of 1m- and 3-month old mice while cluster 13 is mostly composed of cells of 18-, 21-, 24- and 30-month old mice (Figure 4f). Differential gene expression revealed that cluster 10 is enriched for an M2-macrophage gene signature (e.g. Il10, H2-Eb1, H2-Ab1, H2-Aa, Cd74, C1qa, Cxcl16, Hexb, Cd81, C1qb, Cd72) while cluster 13 resembles a M1-proinflammatory macrophage state^50^ (e.g. Hp, Itgal, Spex1, Gngt2) (Extended Data Figure 12f; Supplementary Table 10).

The Tabula Muris Senis is a comprehensive resource for the cell biology community which offers a detailed molecular and cell-type specific portrait of aging. We view such a cell atlas as an essential companion to the genome: the genome provides a blueprint for the organism but does not explain how genes are used in a cell type specific manner or how the usage of genes changes over the lifetime of the organism. The cell atlas provides a deep characterization of phenotype and physiology which can serve as a reference for understanding many aspects of the cell biological changes that mammals undergo during their lifespan.

## Extended Data Figure Legends

**Extended Data Figure 1.**
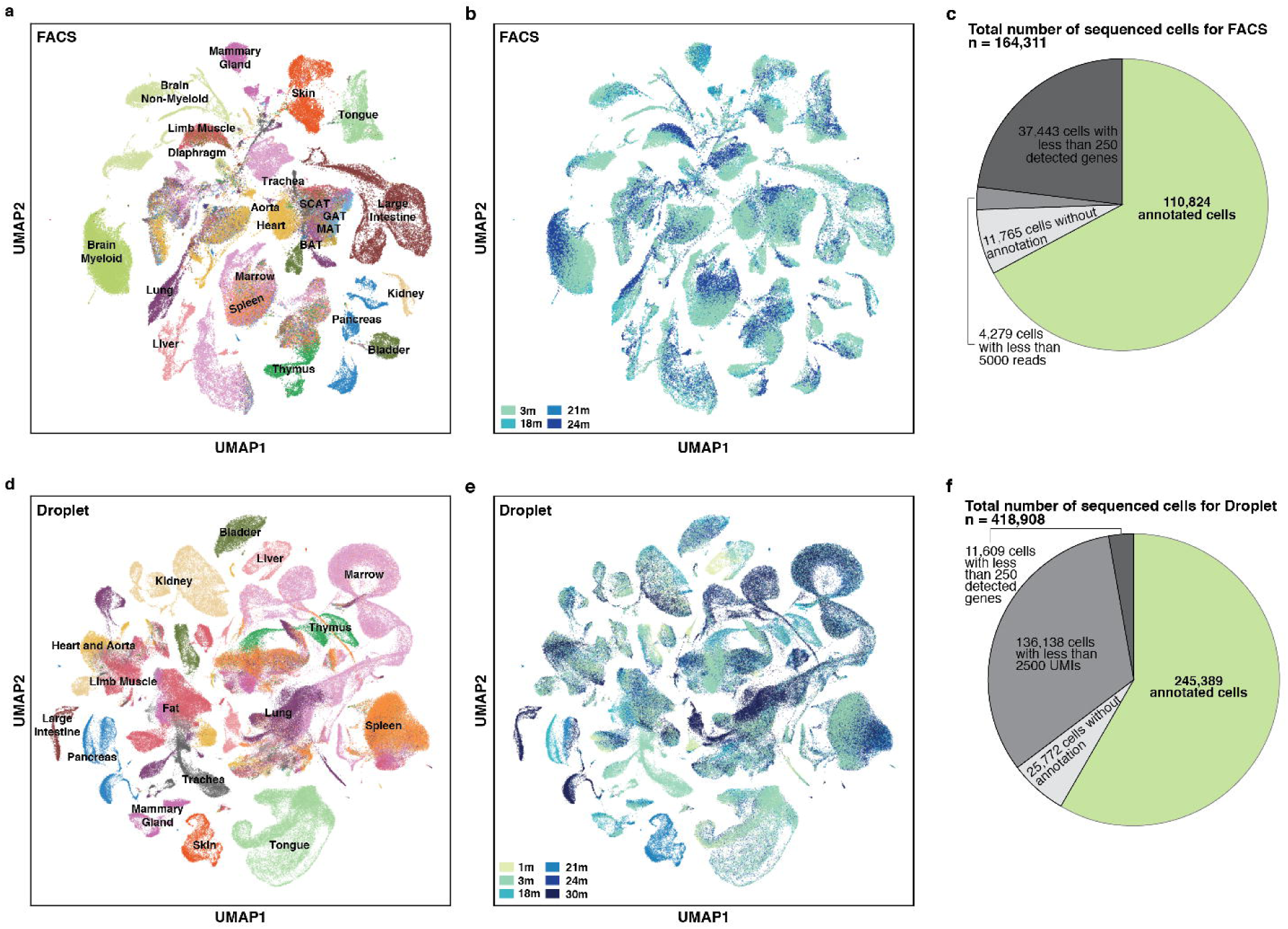
Overview of Tabula Muris Senis (cont.) **a,b**, UMAP plot of all cells collected for FACS colored by tissue (**a**) or age (**b**). **c**, Pie chart with the summary statistics for FACS. **d,e**, UMAP plot of all cells collected for droplet colored by tissue (**d**) or age (**e**). **f**, Pie chart with the summary statistics for droplet.

**Extended Data Figure 2.**
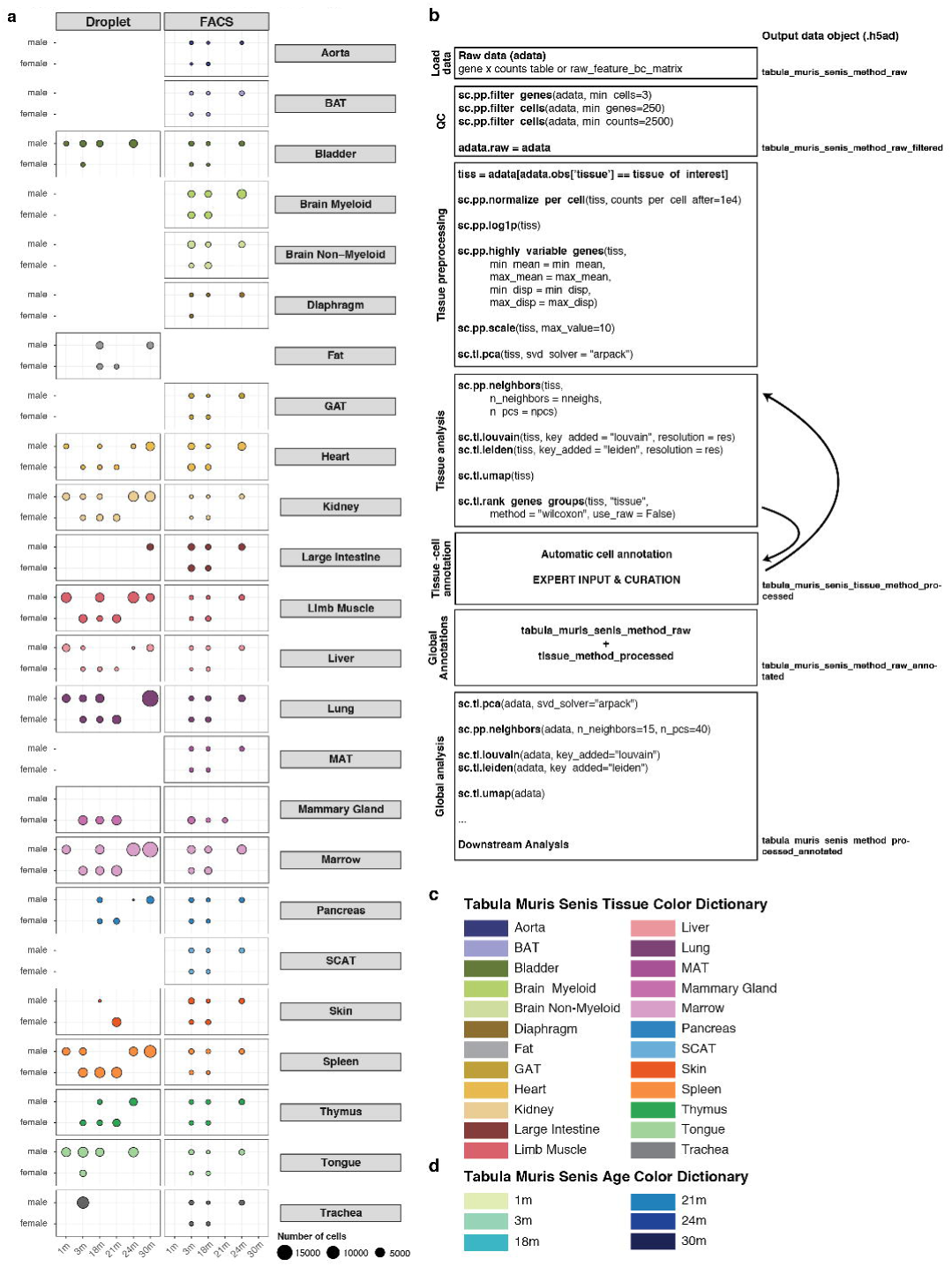
Overview of Tabula Muris Senis (cont.) **a**, Balloon plot showing the number of sequenced cells per sequencing method per organ per sex per age. **b**, Schematic analysis workflow. **c,d,** Tabula Muris Senis color dictionary for organs and tissues (**c**) and ages (**d**).

**Extended Data Figure 3.**
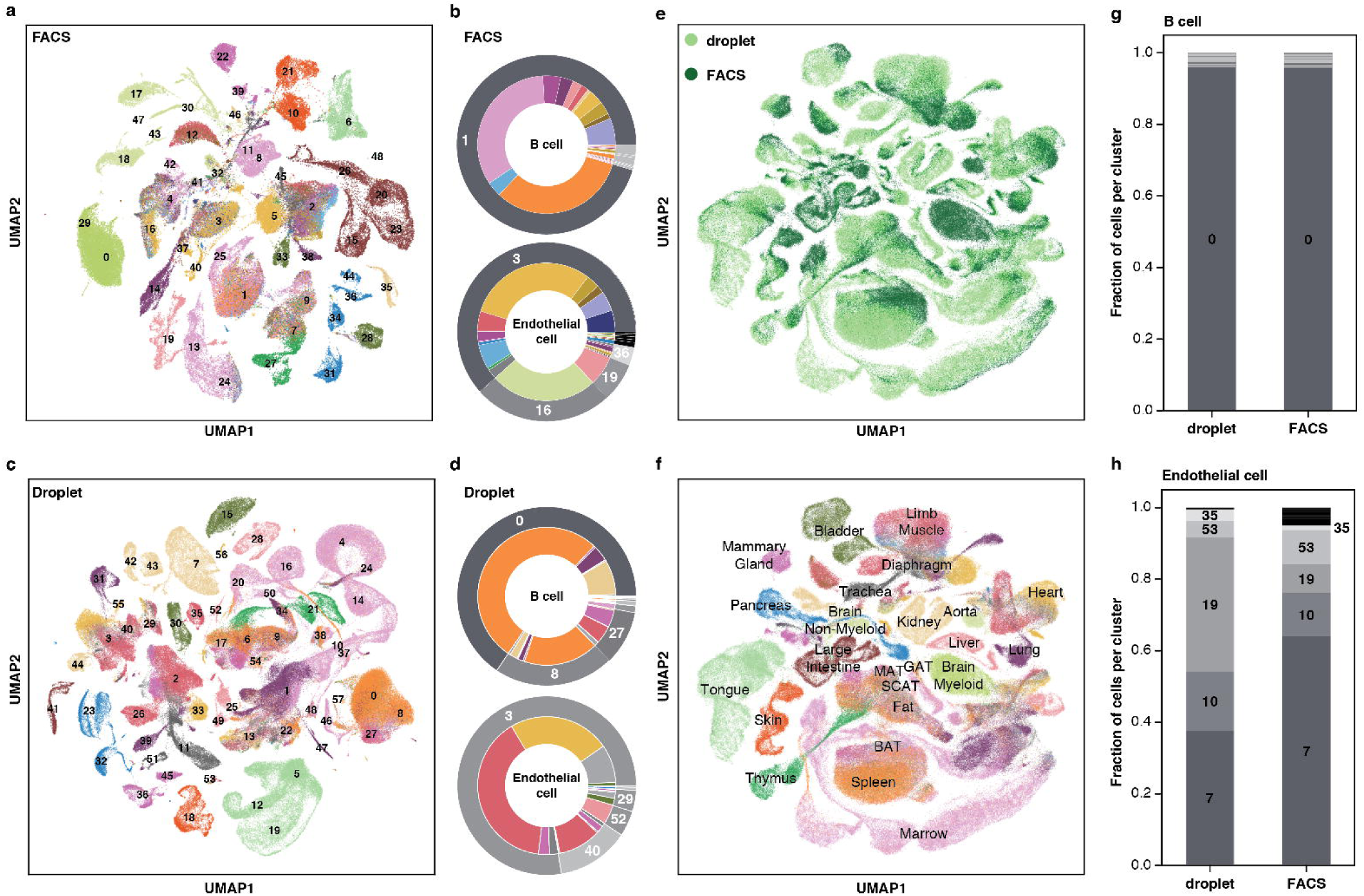
Overview of Tabula Muris Senis (cont.) **a**, UMAP plot of all cells collected by FACS, colored by organ (Extended Data Figure 2c), overlaid with the Louvain cluster numbers; n = 110,824 individual cells. **b**, B cells (top) and endothelial cells (bottom) independently annotated for each organ cluster together by unbiased whole-transcriptome Louvain clustering, irrespectively of the organ they originate from. **c**, UMAP plot of all cells collected by droplet, colored by organ (Extended Data Figure 2c), overlaid with the Louvain cluster numbers; n = 245,389 individual cells. **d**, B cells (and endothelial cells) independently annotated for each organ cluster together by unbiased whole-transcriptome Louvain clustering, irrespectively of the organ where they were found. **e,f**, UMAP plot of all cells collected colored by method (**e**) or tissue (**f**). **g,h**, B cells (**g**) and endothelial cells (**h**) cluster together by unbiased whole-transcriptome Louvain clustering, irrespectively of the technology with which they were found.

**Extended Data Figure 4.**
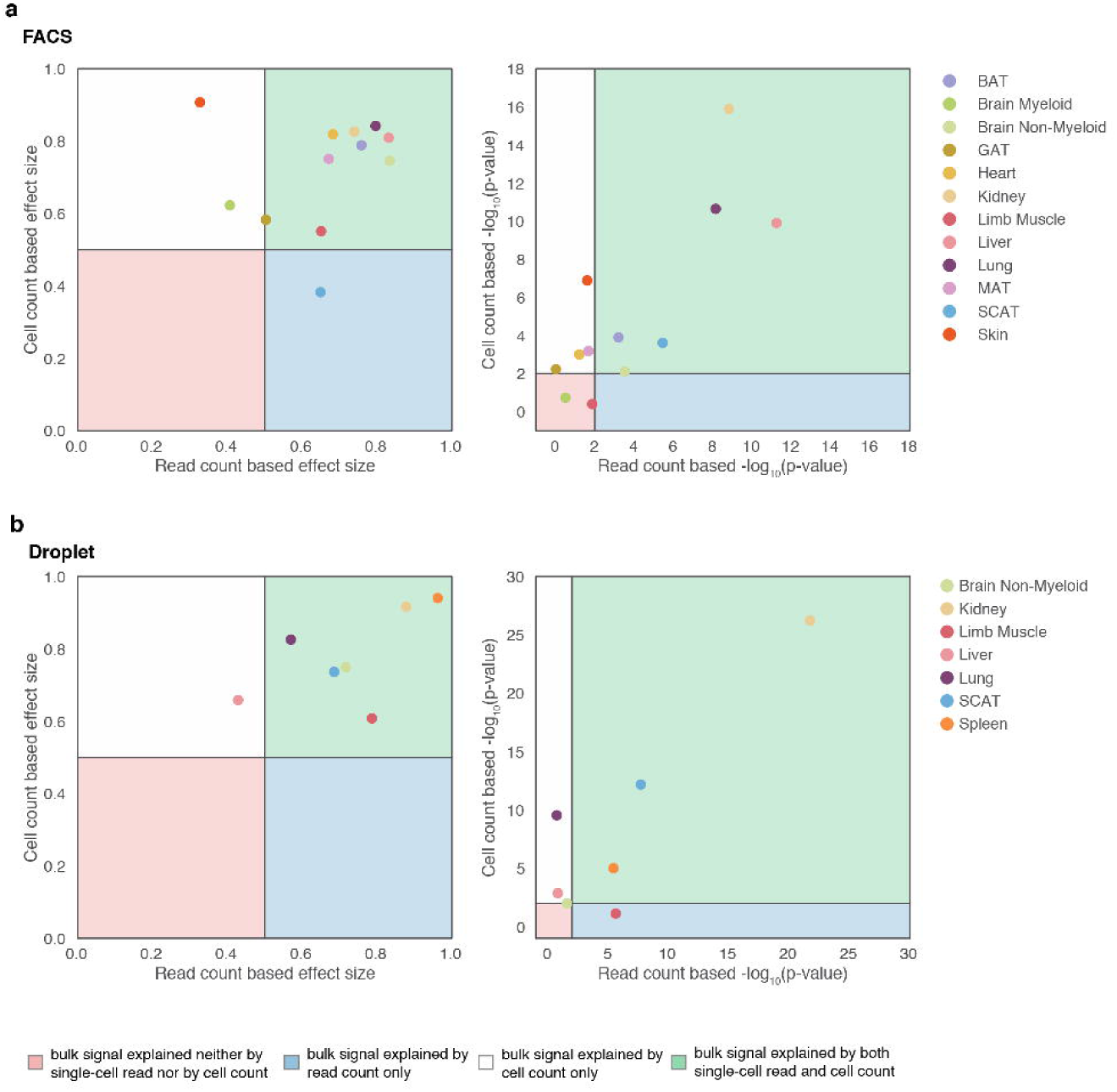
Comparison of bulk and single-cell datasets. Aging patterns from bulk and single-cell data are consistent. Strong changes in bulk gene expression with aging can be either explained by cell or read count-based changes in single-cell data FACS (**a**) and droplet (**b**). Wilcoxon–Mann–Whitney indicates that single-cell data based log_2_ fold-changes of cell or read counts distinguish between up and down regulated genes in bulk data.

**Extended Data Figure 5.**
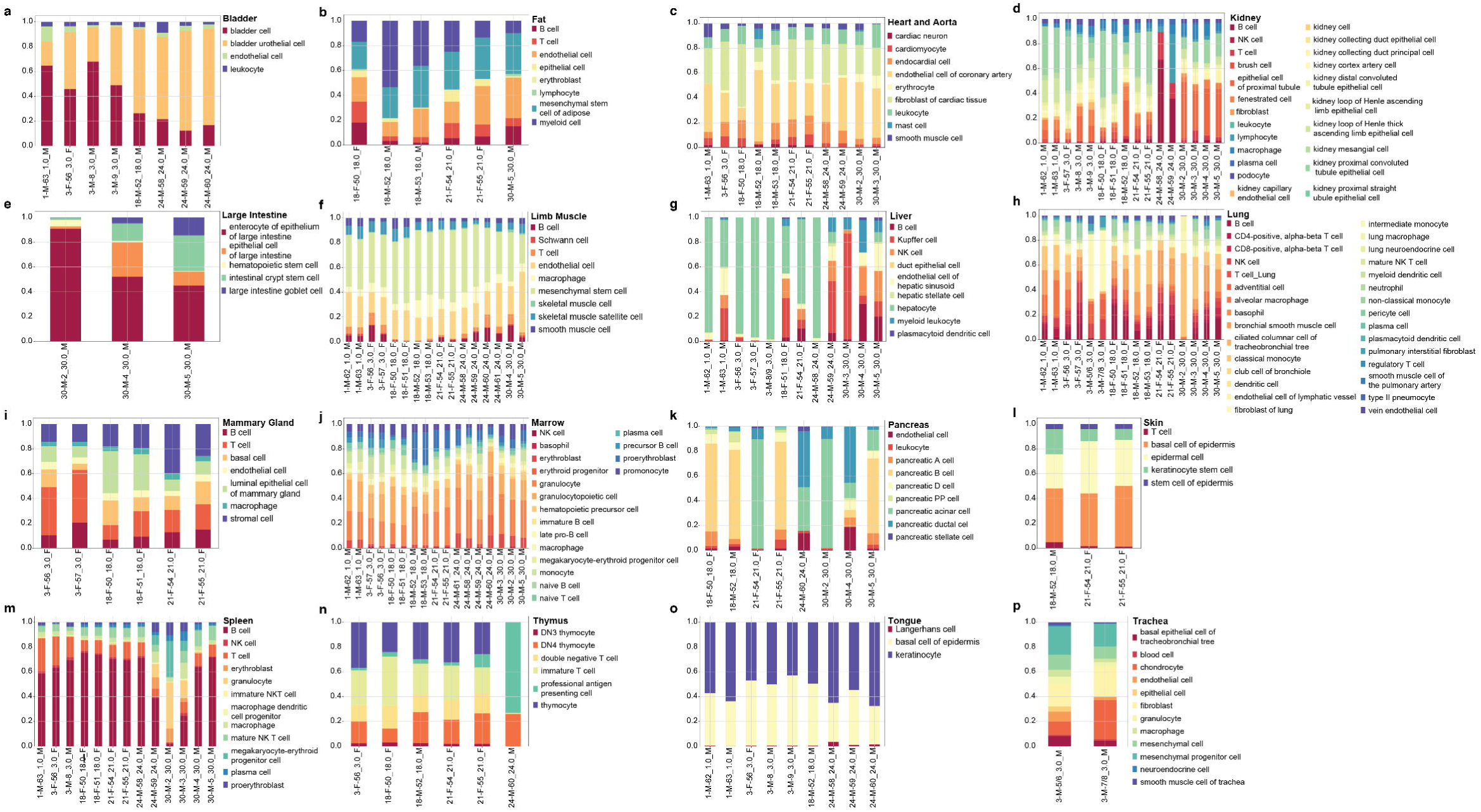
Tissue cell compositions. **a-p**, Alphabetically sorted tissue bar plot showing the relative abundances of cell types in each tissue across the entire age range for the droplet dataset. The tissue cell composition is also available at our online browser <u>tabula-muris-senis.ds.czbiohub.org</u>

**Extended Data Figure 6.**
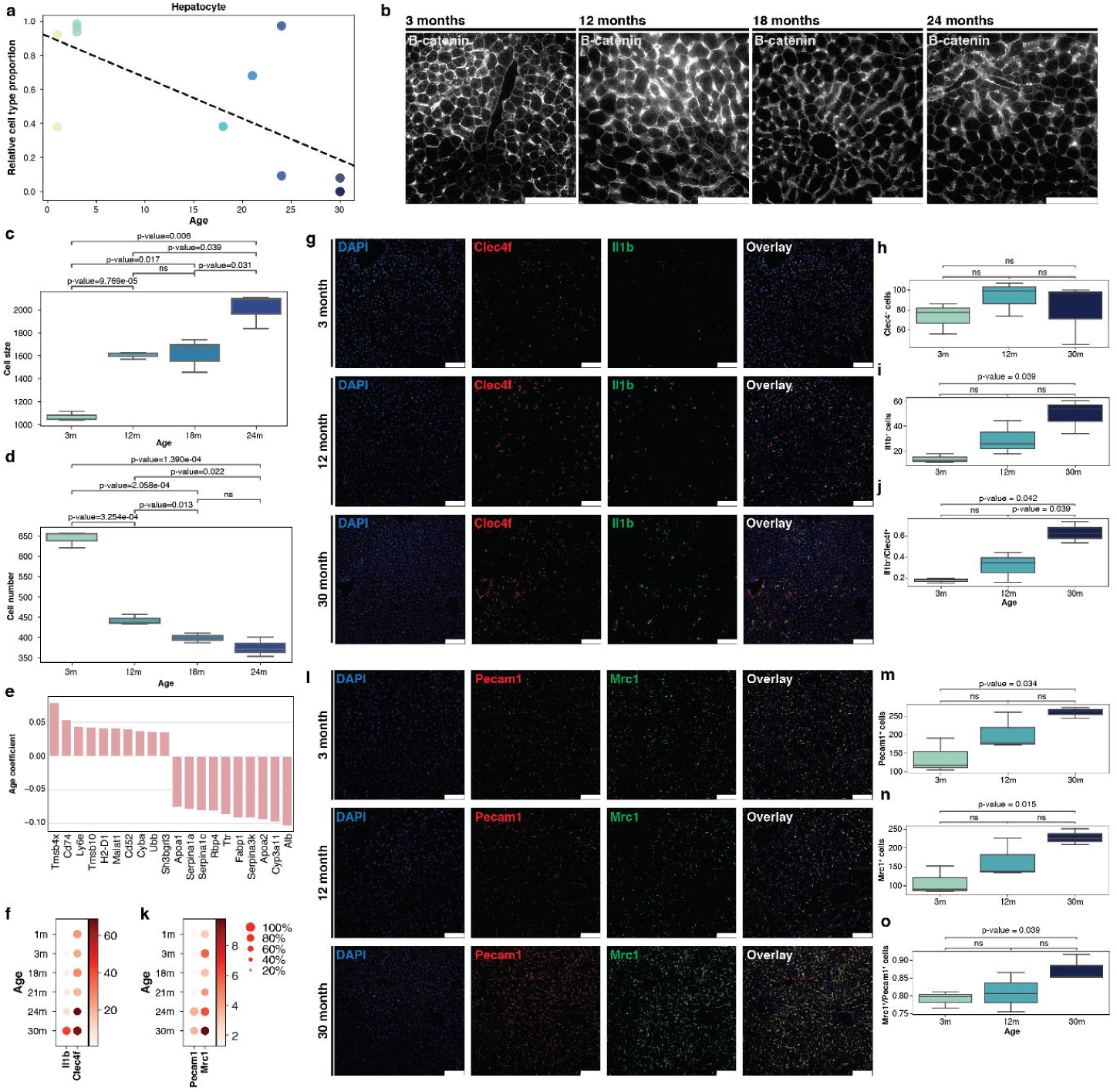
Cellular changes during aging in the liver. **a**, Liver hepatocyte relative abundances change significantly with age (p-value<0.05 and r^2^>0.7 for a hypothesis test whose null hypothesis is that the slope is zero, using two-sided Wald Test with t-distribution of the test statistic). **b-d,** Brightfield imaging of hepatocytes across age (**b**) and respective quantification (**c-d**). **e**, Top 10 upregulated and downregulated genes in liver using age as a continuous covariate while controlling for sex and technology. Genes were classified as significant under an FDR threshold of 0.01 and an age coefficient threshold of 0.005 (corresponding to ∼10% fold change). **f,k,** Gene expression of Il1b and Clec4f (**f**) and Pecam1 and Mrc1 (**k**) in the liver droplet dataset for the six ages. **g-j,** Staining of Kupffer cells across age (**g**) and respective quantification (**h-j**). **l-o,** Staining of liver endothelial cells across ages (**l**) and respective quantification (**m-o**). The white scale bar corresponds to 100µm.

**Extended Data Figure 7.**
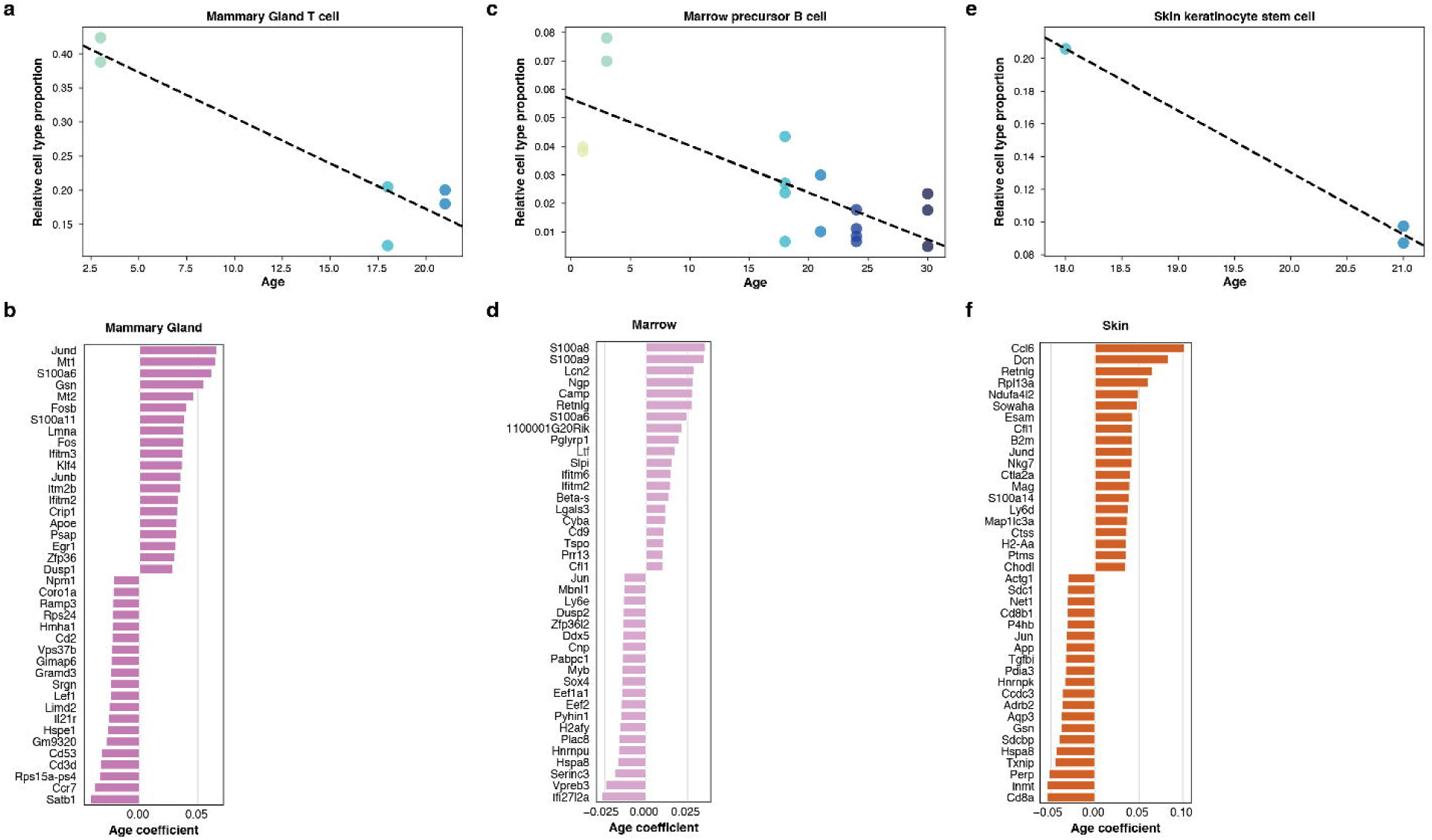
Cellular changes during aging (cont.) **a**, Mammary gland T cell relative abundances change significantly with age (p-value<0.05 and r^2^>0.7 for a hypothesis test whose null hypothesis is that the slope is zero, using two-sided Wald Test with t-distribution of the test statistic). **b,c**, Top 10 upregulated and downregulated genes in mammary gland FACS (**b**) and droplet (**c**) using age as a continuous covariate while controlling for sex. Genes were classified as significant under an FDR threshold of 0.01 and an age coefficient threshold of 0.005 (corresponding to ∼10% fold change). **d**, Marrow precursor B cell relative abundances change significantly with age (p-value<0.05 and r^2^>0.7 for a hypothesis test whose null hypothesis is that the slope is zero, using two-sided Wald Test with t-distribution of the test statistic). **e,f**, Top 10 upregulated and downregulated genes in marrow FACS (**e**) and droplet (**f**) using age as a continuous covariate while controlling for sex. Genes were classified as significant under an FDR threshold of 0.01 and an age coefficient threshold of 0.005 (corresponding to ∼10% fold change). **g**, Skin keratinocyte stem cell relative abundances change significantly with age (p-value<0.05 and r^2^>0.7 for a hypothesis test whose null hypothesis is that the slope is zero, using two-sided Wald Test with t-distribution of the test statistic). **h**, Top 10 upregulated and downregulated genes in skin FACS using age as a continuous covariate while controlling for sex. Genes were classified as significant under an FDR threshold of 0.01 and an age coefficient threshold of 0.005 (corresponding to ∼10% fold change).

**Extended Data Figure 8.**
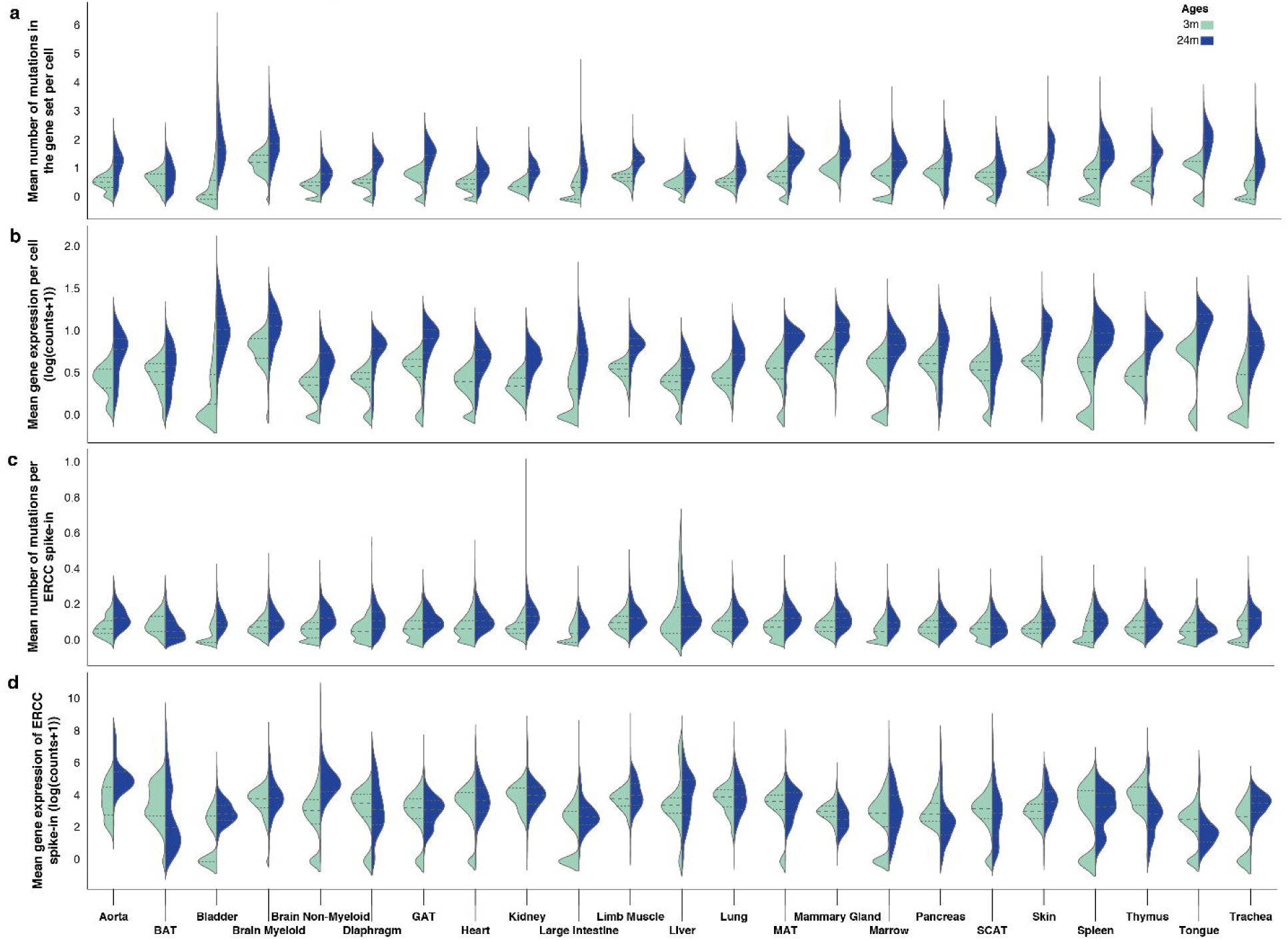
Mutational burden across tissues in the aging mice (cont. 24m vs 3m). **a,b**, Mean number of somatic mutations (**a**) and raw expression (**b**) across all tissues per age group (3m and 24m). **c,d**, Mean number of mutations in ERCC spike-in (**c**) and ERCC raw expression (**d**) across all tissues per age group (3m and 24m). Mutations are presented as the mean number of mutations per gene per cell.

**Extended Data Figure 9.**
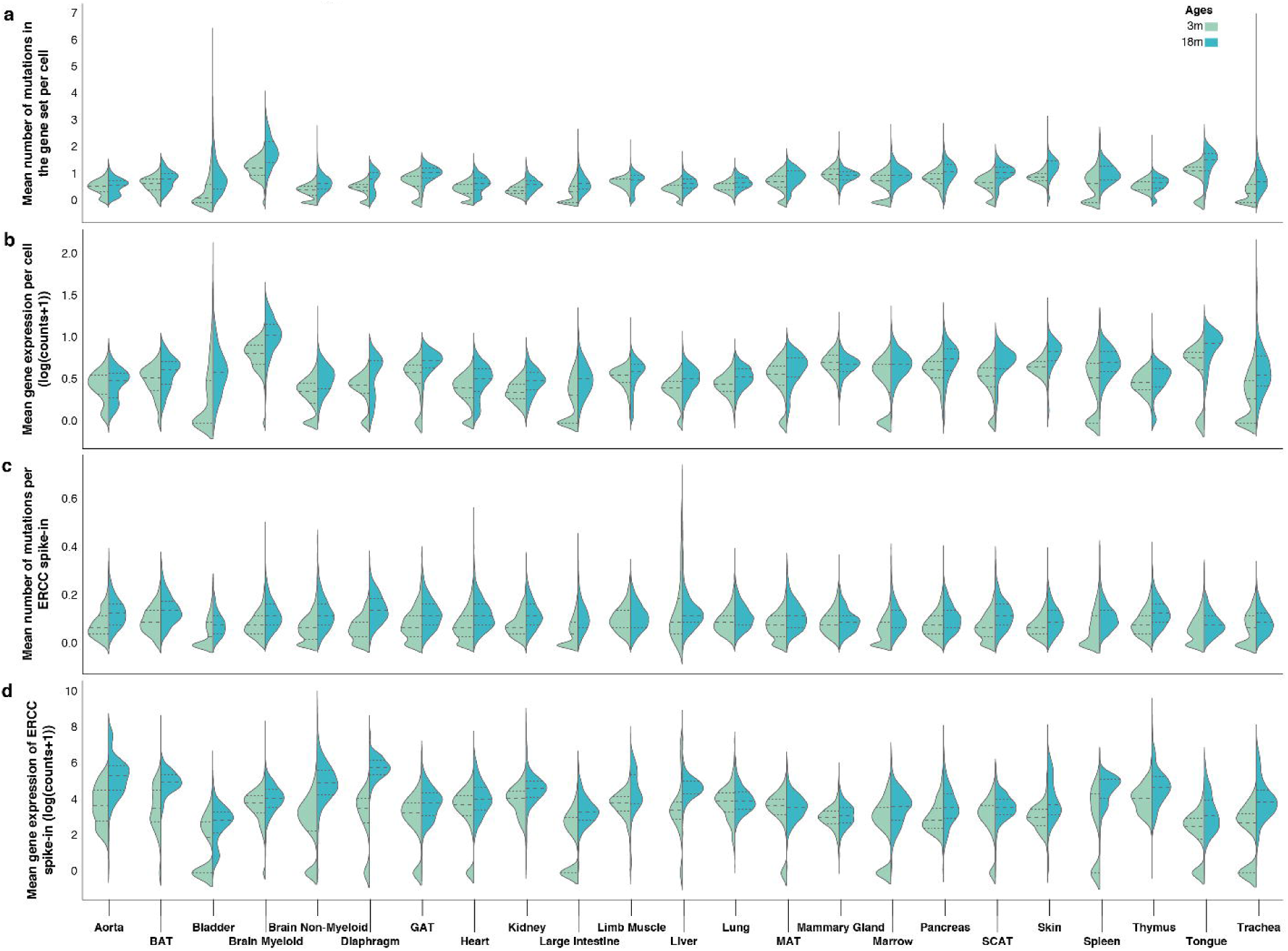
Mutational burden across tissues in the aging mice (cont. 18m vs 3m). **a,b**, Mean number of somatic mutations (**a**) and raw expression (**b**) across all tissues per age group (3m and 18m). **c,d**, Mean number of mutations in ERCC spike-in (**c**) and ERCC raw expression (**d**) across all tissues per age group (3m and 18m). Mutations are presented as the mean number of mutations per gene per cell.

**Extended Data Figure 10.**
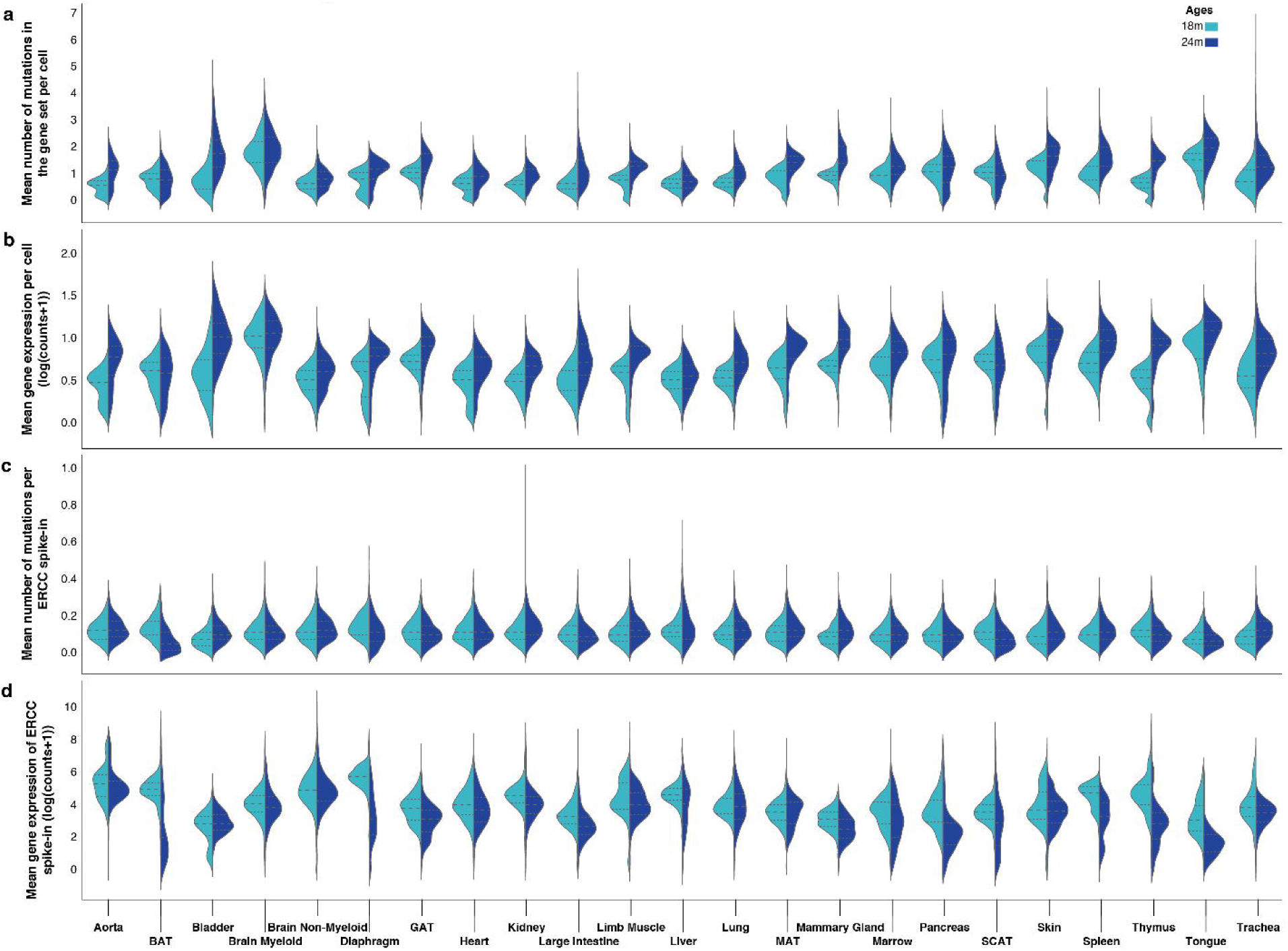
Mutational burden across tissues in the aging mice (cont. 24m vs 18m). **a,b**, Mean number of somatic mutations (**a**) and raw expression (**b**) across all tissues per age group (18m and 24m). **c,d**, Mean number of mutations in ERCC spike-in (**c**) and ERCC raw expression (**d**) across all tissues per age group (18m and 24m). Mutations are presented as the mean number of mutations per gene per cell.

**Extended Data Figure 11.**
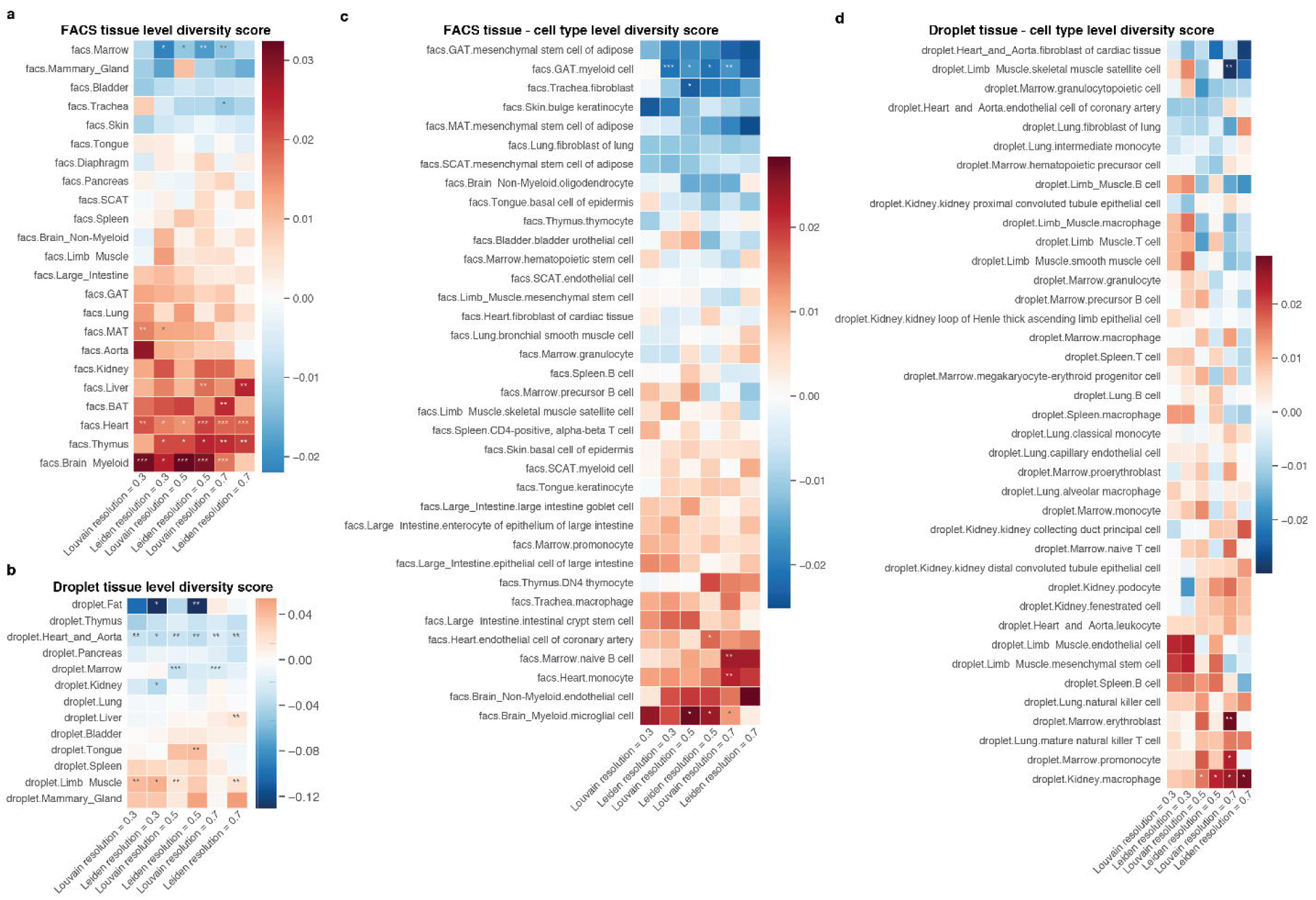
Diversity score summary. **a,b**, Heatmap summary of the overall tissue diversity score for FACS (**a**) and droplet (**b**). **c,d**, Heatmap summary of the tissue cell-type diversity score for FACS (**c**) and droplet (**d**).

**Extended Data Figure 12.**
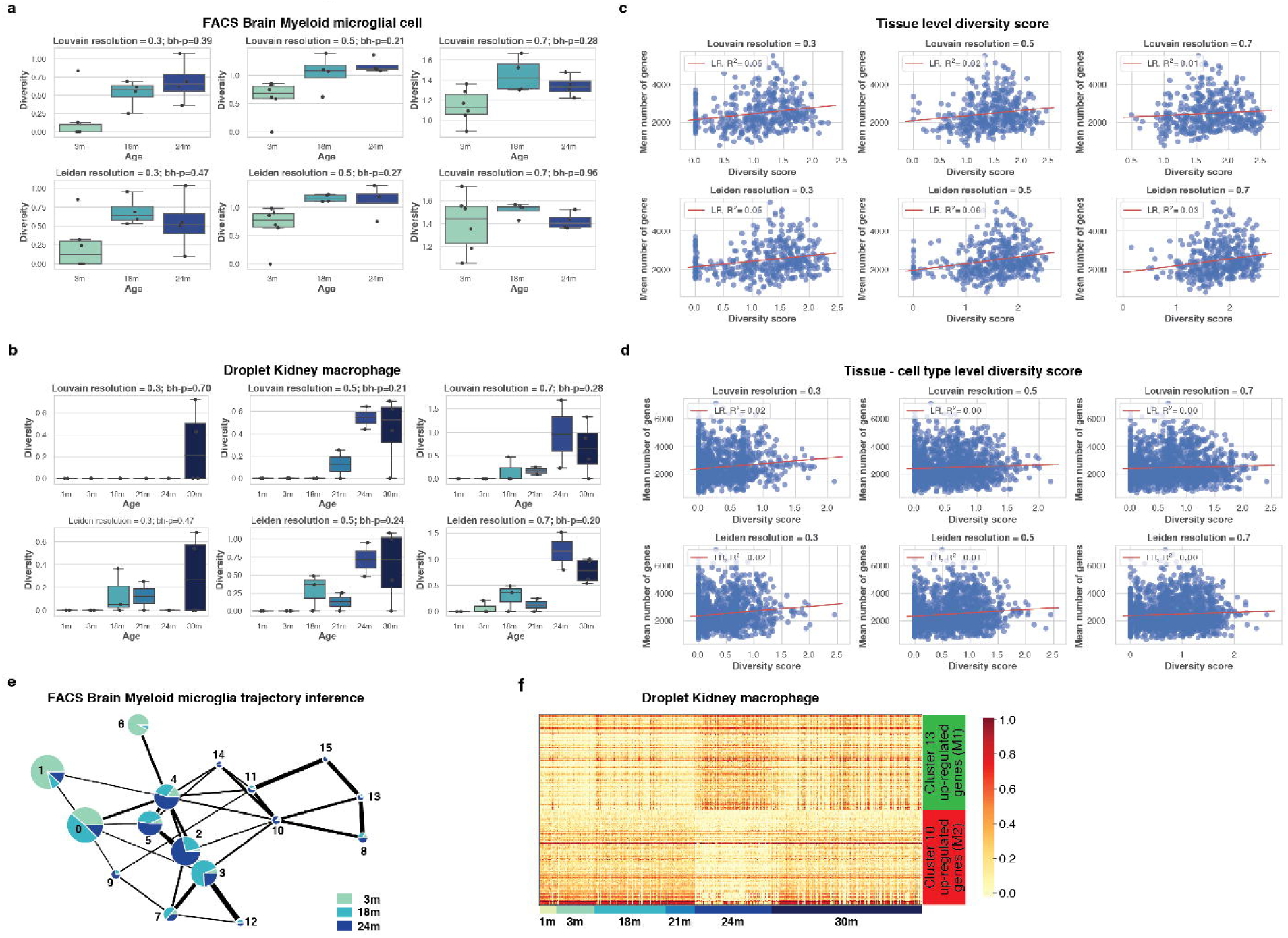
The aging immune system (cont.) **a,b,** Diversity score at different cluster resolutions for FACS brain myeloid microglia cell (**a**) and droplet kidney macrophage (**b)**. **c,d**, Diversity score correlation with the number of genes expressed per tissue (**c**) or tissue cell-type (**d**). **e**, PAGA^51^ trajectory for brain myeloid microglia cell. **f**, Differential gene expression analysis of cluster 10 (mostly young macrophages) versus clusters 13 (mostly old macrophages). For the complete gene list please refer to Supplementary Table 10.

## Supplementary Figure Legends

**Supplementary Figure 1.**
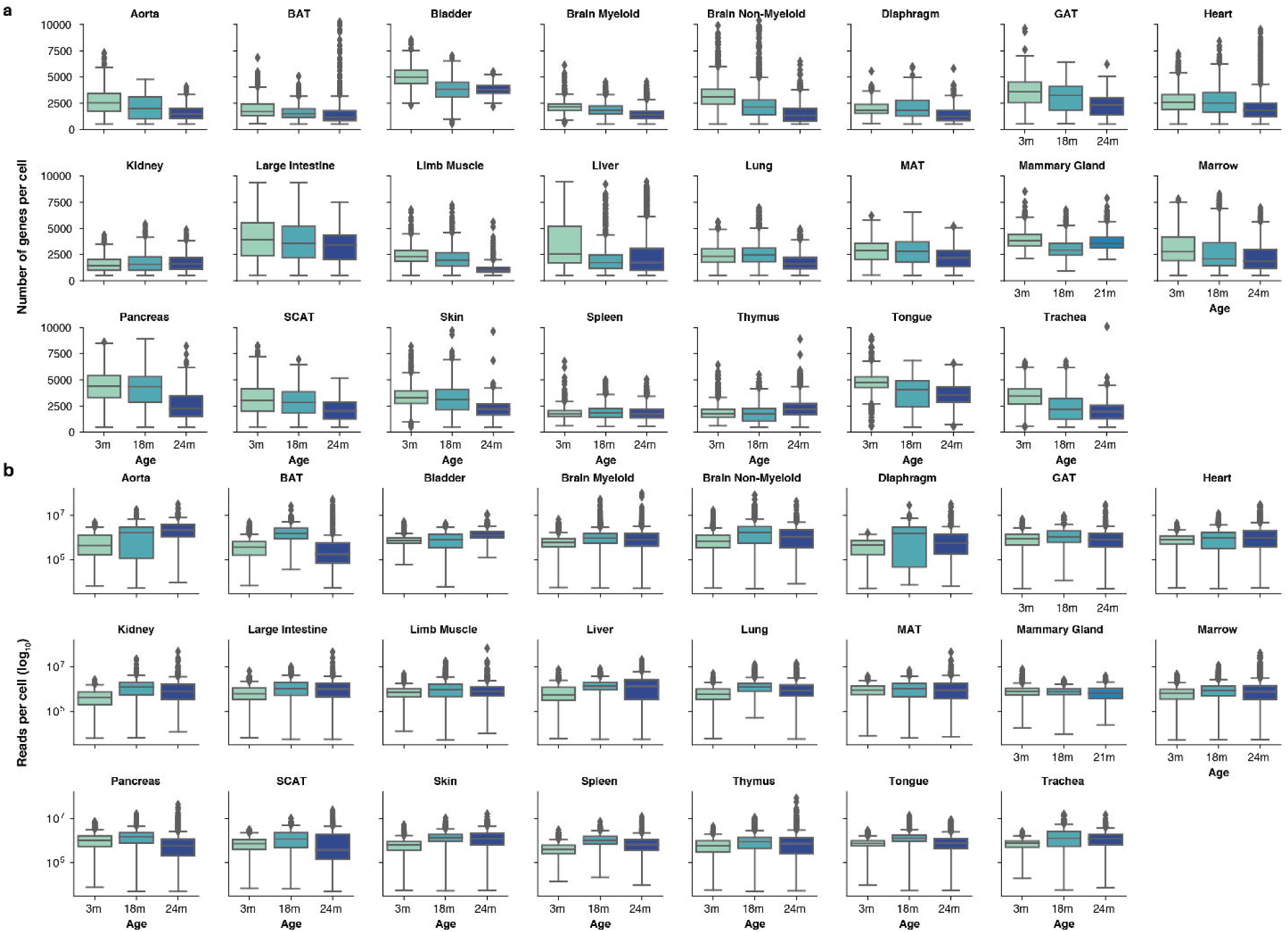
FACS sequencing statistics. **a**, Box plot of the number of genes detected per cell for each organ and age. **b**, Box plot of the number of reads per cell (log-scale) for each organ and age.

**Supplementary Figure 2.**
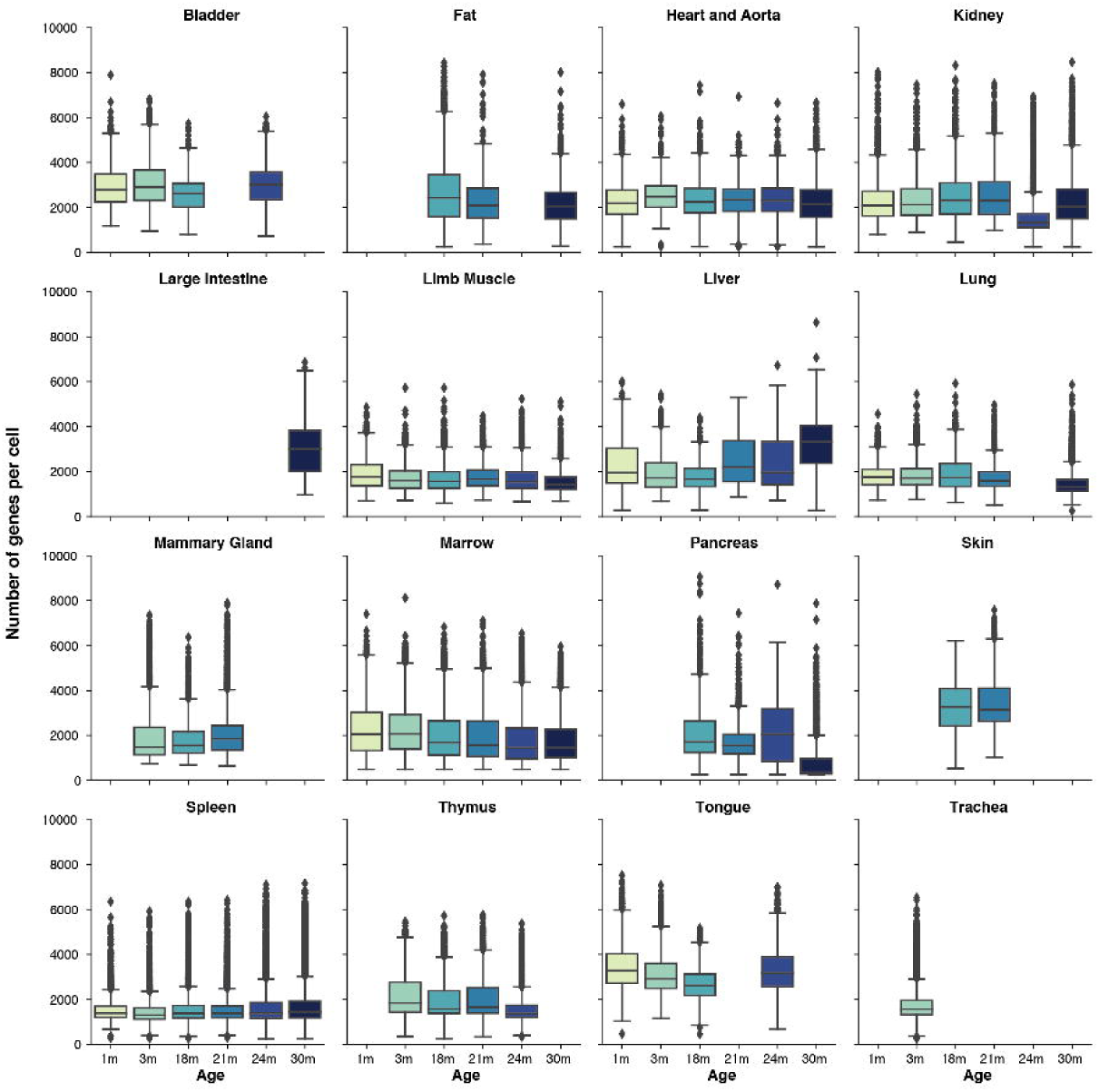
Droplet sequencing statistics. Box plot of the number of genes detected per cell for each organ and age.

**Supplementary Figure 3.**
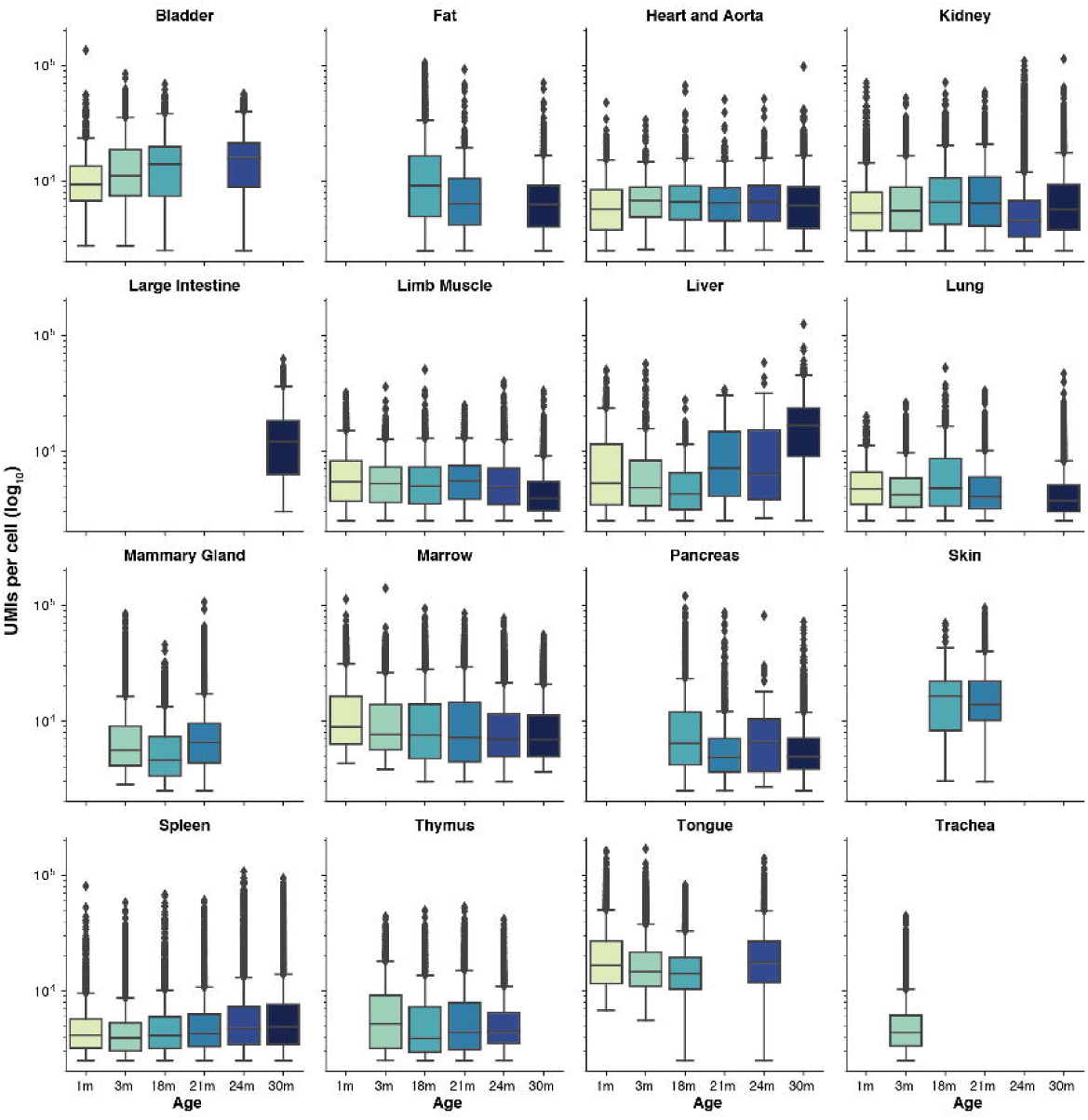
Droplet sequencing statistics (cont.) Box plot of the number of UMIs per cell (log-scale) for each organ and age.

## Supplementary Tables

**Supplementary Table 1. Summary of the FACS dataset.**

**a**, Number of cells grouped by age, sex, mouse id and tissue. **b**, Number of cells grouped by tissue, cell ontology class and age. **c**, Number of cells grouped by Louvain cluster number, cell ontology class, tissue and age. **d**, Number of cells grouped by cell ontology class, Louvain cluster number, tissue and age. **e**, Fraction of cells in each Louvain cluster per cell ontology class and tissue. **f**, Fraction of cells in each Louvain cluster per tissue. **g**, Fraction of cells in each Louvain cluster per cell ontology class.

**Supplementary Table 2. Summary of the droplet dataset.**

**a**, Number of cells grouped by age, sex, mouse id and tissue. **b**, Number of cells grouped by tissue, cell ontology class and age. **c**, Number of cells grouped by Louvain cluster number, cell ontology class, tissue and age. **d**, Number of cells grouped by cell ontology class, Louvain cluster number, tissue and age. **e**, Fraction of cells in each Louvain cluster per cell ontology class and tissue. **f**, Fraction of cells in each Louvain cluster per tissue. **g**, Fraction of cells in each Louvain cluster per cell ontology class.

**Supplementary Table 3. Summary of the batch corrected dataset.**

**a**, Number of cells grouped by age, sex, mouse id and tissue. **b**, Number of cells grouped by tissue, cell ontology class and age. **c**, Number of cells grouped by Louvain cluster number, cell ontology class, tissue and age. **d**, Number of cells grouped by cell ontology class, Louvain cluster number, tissue and age. **e**, Fraction of cells in each Louvain cluster per cell ontology class and tissue. **f**, Fraction of cells in each Louvain cluster per tissue. **g**, Fraction of cells in each Louvain cluster per cell ontology class.

**Supplementary Table 4. Cellular fraction changes for senescence markers.** This supplementary table supports Figure 2a-d.

**Supplementary Table 5. Cellular fraction changes.** This supplementary table supports Figure 2e,g,i; Extended Data Figure 5; Extended Data Figure 6a and Extended Data Figure 7a,c,e.

**Supplementary Table 6. Differential gene expression analysis.** This supplementary table supports Figure 2f,h,j; Extended Data Figure 6e and Extended Data Figure 7b,d,f.

**Supplementary Table 7. Quantification of Liver in-situ staining.** This supplementary table supports Figure 6b-d,g-j and l-o. fov stands for field of view.

**Supplementary Table 8. Summary statistics for the GATK analysis. Cell** is the unique cell identifier; **ercc** is the average number of mutations per cell found in the ERCC spike-in, **adata** is the average number of mutations per cell in the gene set of the tissue; **ercc_raw_counts** are the average number of ERCC spike-in counts per cell and **ercc_counts** are the log(ercc_raw_counts+1); **adata_raw_counts** are the average number of gene counts per cell and **adata_counts** are the log(adata_raw_counts+1); **tissue**, **age** and **cell_ontology_class** are the metadata of the respective cell id and **agenum** is the age as a numerical variable; **functional_annotations** is a categorical variable binning each cell type as endothelial, immune, parenchymal, stem cell/progenitor or stromal.

**Supplementary Table 9. B-cell and T-cell repertoire analysis raw data.** This table supports Figure 4a,b.

**Supplementary Table 10. Differential gene expression for the tissue cell type whose diversity significantly changes with age. a**, FACS brain myeloid microglia differentially upregulated genes between clusters 10, 12 and 14 versus clusters 1 and 6**. b**, FACS brain myeloid microglia differentially upregulated genes between clusters 1 and 6 versus clusters 10, 12 and 14**. c**, Droplet kidney macrophage differentially upregulated genes between cluster 13 and cluster 10**. d**, Droplet kidney macrophage differentially upregulated genes between cluster 10 and cluster 13**. e,** Alzheimer’s disease microglia signature from^47^. This table supports Figure 4d,e and Extended Data Figure 12f.

## Methods

All data, protocols, analysis scripts and an interactive data browser are publicly available.

### Experimental Procedures

#### Mice and organ collection

Male and virgin female C57BL/6JN mice were shipped from the National Institute on Aging colony at Charles River (housed at 67–73 °F) to the Veterinary Medical Unit (VMU; housed at 68–76 °F)) at the VA Palo Alto (VA). At both locations, mice were housed on a 12-h light/dark cycle and provided food and water *ad libitum*. The diet at Charles River was NIH-31, and Teklad 2918 at the VA VMU. Littermates were not recorded or tracked, and mice were housed at the VA VMU for no longer than 2 weeks before euthanasia, with the exception of mice older than 18 months, which were housed at the VA VMU beginning at 18 months of age. Before tissue collection, mice were placed in sterile collection chambers at 8 am for 15 min to collect fresh fecal pellets. After anaesthetization with 2.5% v/v Avertin, mice were weighed, shaved, and blood was drawn via cardiac puncture before transcardial perfusion with 20 ml PBS. Mesenteric adipose tissue was then immediately collected to avoid exposure to the liver and pancreas perfusate, which negatively affects cell sorting. Isolating viable single cells from both the pancreas and the liver of the same mouse was not possible; therefore, two males and two females were used for each. Whole organs were then dissected in the following order: large intestine, spleen, thymus, trachea, tongue, brain, heart, lung, kidney, gonadal adipose tissue, bladder, diaphragm, limb muscle (tibialis anterior), skin (dorsal), subcutaneous adipose tissue (inguinal pad), mammary glands (fat pads 2, 3 and 4), brown adipose tissue (interscapular pad), aorta and bone marrow (spine and limb bones). Organ collection concluded by 10 am. After single-cell dissociation as described below, cell suspensions were either used for FACS of individual cells into 384-well plates, or for preparation of the microfluidic droplet library. All animal care and procedures were carried out in accordance with institutional guidelines approved by the VA Palo Alto Committee on Animal Research.

#### Tissue dissociation and sample preparation

All tissues were processed as previously described^5^.

#### Sample size, randomization and blinding

No sample size choice was performed before the study. Randomization and blinding were not performed: the authors were aware of all data and metadata-related variables during the entire course of the study.

#### Single-cell methods

All protocols used in this study are described in detail elsewhere^5^. Those include: i) preparation of lysis plates, ii) FACS sorting, iii) cDNA synthesis using the Smart-seq2 protocol^52, 53^, iv) library preparation using an in-house version of Tn5^54, 55^,v) library pooling and Quality control and vi) sequencing. For further details please refer to http://dx.doi.org/10.17504/protocols.io.2uwgexe

#### Microfluidic droplet single-cell analysis

Single cells were captured in droplet emulsions using the GemCode Single-Cell Instrument (10x Genomics) and scRNA-seq libraries were constructed as per the 10x Genomics protocol using GemCode Single-Cell 3’ Gel Bead and Library V2 Kit. In brief, single cell suspensions were examined using an inverted microscope, and if sample quality was deemed satisfactory, the sample was diluted in PBS with 2% FBS to a concentration of 1000 cells per μl. If cell suspensions contained cell aggregates or debris, two additional washes in PBS with 2% FBS at 300*g*for 5 min at 4 °C were performed. Cell concentration was measured either with a Moxi GO II (Orflo Technologies) or a haemocytometer. Cells were loaded in each channel with a target output of 5,000 cells per sample. All reactions were performed in the Biorad C1000 Touch Thermal cycler with 96-Deep Well Reaction Module. 12 cycles were used for cDNA amplification and sample index PCR. Amplified cDNA and final libraries were evaluated on a Fragment Analyzer using a High Sensitivity NGS Analysis Kit (Advanced Analytical). The average fragment length of 10x cDNA libraries was quantitated on a Fragment Analyzer (AATI), and by qPCR with the Kapa Library Quantification kit for Illumina. Each library was diluted to 2 nM, and equal volumes of 16 libraries were pooled for each NovaSeq sequencing run. Pools were sequenced with 100 cycle run kits with 26 bases for Read 1, 8 bases for Index 1, and 90 bases for Read 2 (Illumina 20012862). A PhiX control library was spiked in at 0.2 to 1%. Libraries were sequenced on the NovaSeq 6000 Sequencing System (Illumina).

### Computational methods

#### Data extraction

Sequences from the NovaSeq were de-multiplexed using bcl2fastq version 2.19.0.316. Reads were aligned using to the mm10plus genome using STAR version 2.5.2b with parameters TK. Gene counts were produced using HTSEQ version 0.6.1p1 with default parameters, except ‘stranded’ was set to ‘false’, and ‘mode’ was set to ‘intersection-nonempty’. Sequences from the microfluidic droplet platform were de-multiplexed and aligned using CellRanger version 2.0.1, available from 10x Genomics with default parameters.

#### Data pre-processing

Gene count tables were combined with the metadata variables using the Scanpy^56^ Python package version 1.4. We removed genes not expressed in at least 3 cells and then cells that did not have at least 250 detected genes. For FACS we removed cells with less than 5000 counts and for droplet cells with less than 2500 UMIs. The data was then normalized using size factor normalization such that every cell has 10,000 counts and log transformed. We computed highly variable genes using default parameters and then scaled the data to a maximum value of 10. After we computed PCA, neighborhood graph and clustered the data using Louvain^7^ and Leiden^8^ methods. The data was visualized using UMAP projection. When performing batch correction to remove the technical artifacts introduced by the technologies, we replaced the neighborhood graph computation with **bbknn**^6^. Step-by-step instructions to reproduce the pre-processing of the data are available from GitHub.

#### Cell type annotation

To define cell types we analyzed each organ independently but combining all ages. In a nutshell, we performed principal component analysis on the most variable genes between cells, followed by Louvain and Leiden graph-based clustering. Next we subset the data for 3m (Tabula Muris^5^) and compute how many cell types map to each individual cluster. For the clusters that we had a single 1:1 mapping (cluster:cell type) we propagate the annotations for all ages; in case there is a 1:many mapping we flagged that cluster for manual validation. Step-by-step instructions to reproduce this method are available from GitHub. For each cluster, we provide annotations in the controlled vocabulary of the cell ontology^57^ to facilitate inter-experiment comparisons. Using this method, we were able to annotate automatically (∼1min per tissue) over 70% of the dataset. The automatic annotations were then reviewed by each of the tissue experts leading to a fully curated dataset for all the cell types in Tabula Muris Senis.

#### Tissue cell composition analysis

For each tissue and age, we computed the relative proportion of each cell type. Next we used scipy.stats linregress to regress the relative tissue-cell type changes against age and considered significant the changes with p-value<0.05 for a hypothesis test whose null hypothesis is that the slope is zero, using two-sided Wald Test with t-distribution of the test statistic and a r^2^>0.5.

#### Differential gene expression

We performed differential gene expression analysis on each tissue with a well-powered sample size (>100 cells in both young (1m and 3m) and old age group (18m, 21m, 24m and 30m)). We use a linear model^58^ treating age as a numerical variable while controlling for sex and technology. We apply a false-discovery rate (FDR) threshold of 0.01 and an age coefficient threshold of 0.005 (corresponding to ∼10% fold change).

#### In Situ RNA Hybridization and quantification

*In situ* RNA hybridization was performed using the Advanced Cell Diagnostics RNAscope® Multiplex Fluorescent Detection kit v2 (323110, Bio-techne) according to the manufacturer’s instructions. Staining of mouse liver specimens was performed using 5μm paraffin-embedded thick sessions. Mouse livers were fixed in 10% formalin buffer saline (HT501128, Sigma Aldrich) for 24h at room temperature before paraffin embedding. For multiplex staining the following probes were used; *Clec4f* (Mm-Clec4f 480421, *Il1b* (Mm-Il1b 316891-C2), *Pecam1* (Mm-Pecam-1 316721), *Mrc1* (Mm-Mrc1 437511-C3). Slides were counter stained with Prolong gold antifade reagent with DAPI (P36931, Life technologies). Mounted slides were imaged on a Leica DM6 B fluorescent microscope (Leica Biosystems). Image quantification was performed using the starfish open source image-based transcriptomics pipeline (please refer to Starfish: Open Source Image Based Transcriptomics and Proteomics Tools available from http://github.com/spacetx/starfish and ^59^).

#### Comparison between bulk and single-cell datasets

The differential gene analysis was defined on a per tissue basis. First, we investigated genes based on the single-cell data. We only considered cells from male animals and perform our analysis on the log (1 + CPM) transformed single-cell count matrices. Note that normalization of the single-cell data was done on a per cell basis. We defined two group of cells based on age: young cells with age <= 3 months (Y) and old cells with age > 3 months (O). For each gene we compute the log_2_ fold-change of cell and read counts between O and Y. We defined cell count as the fraction of cells that express the gene. Similarly, we defined read count as the mean read count of the gene in the cells that express it. The calculated log_2_ fold-changes of a gene reflect its expression changes with aging within the single-cell data. Next we analyze each gene based on the bulk data. We computed the Spearman (Sp) correlation of bulk DESeq2 normalized gene expression with aging. We defined two groups of genes based on the bulk data, increasing with age Sp > 0.7 (U) and decreasing with age Sp < −0.7 (D). Finally, we compared the single-cell data based log_2_ fold-changes between the bulk data defined groups U and D. Specifically, we run Wilcoxon–Mann–Whitney test in order to understand if log_2_ fold-changes of cell or read counts could distinguish between the two groups. We used the U statistic for effect size.

#### T-Cell processing

We used TraCeR^44^ version 0.5 to identify T-Cell clonal populations. We ran tracer assemble with --species Mmus set. We then ran tracer summarise with –species Mmus to create the final results. We used the following versions for TraCeR dependencies: igblast version 1.7.0, kallisto version v0.43.1, Salmon version 0.8.2, Trinity version v2.4.0, GRCm38 reference genome. Step-by-step instructions to reproduce the processing of the data are available from GitHub.

#### B-Cell processing

We used singlecell-ige^43^ version eafb6d126cc2d6511faae3efbd442abd7c6dc8ef (https://github.com/dcroote/singlecell-ige) to identify B-Cell clonal populations. We used the default configuration settings except we set the species to mouse. Step-by-step instructions to reproduce the processing of the data are available from GitHub.

#### Mutation analysis

We used samtools^60^ version 1.9 and GATK^39^ version v4.1.1.0 for mutation analysis. We used samtools faidx to create our index file. Then we used GATK CreateSequenceDictionary and GRCm38, as the reference, to create our sequence dictionary. Next we used GATK AddOrReplaceReadGroups to create a single read group using parameters -RGID 4 -RGLB lib1 -RGPL illumina-RGPU unit1 –RGSM 20. Finally we used GATK HaplotypeCaller to call the mutations. We disabled the following read filters: MappingQualityReadFilter, GoodCigarReadFilter, NotSecondaryAlignmentReadFilter, MappedReadFilter, MappingQualityAvailableReadFilter, NonZeroReferenceLengthAlignmentReadFilter, NotDuplicateReadFilter, PassesVendorQualityCheckReadFilter, and WellformedReadFilter, but kept all other default settings. The results were summarized per gene in the form of a mutation count per cell table. We started by removing genes mutated in over 60% of cells, to eliminate the possible bias of germline mutations. Then for each tissue we selected genes expressed in at least 75% of the cells for all the time points to avoid confounding the mutation results with differential gene expression associated with age. Next we computed the average number of mutations in the gene set (or ERCC spike-in controls) per cell and also the average number of raw counts (Supplementary Table 8) and plotted the different distributions. Step-by-step instructions to reproduce the processing of the data are available from GitHub.

#### Diversity score

The raw FACS or droplet dataset were used as the input. We filtered genes expressed in fewer than 5 cells, filtered cells if expressing fewer than 500 genes and discarded cells with total number of counts less than 5000. Next we performed size factor normalization such that every cell had 1e4 counts and performed a log1p transformation. This was followed by clustering, where we clustered every tissue and every tissue-cell type for every mouse separately using 6 different configurations: resolution parameters (0.3, 0.5, 0.7) * clustering method (Louvain, Leiden). This is to provide a robust clustering result. For each combination (each tissue-mouse and each tissue-cell_type-mouse), we computed the clustering diversity score as the Shannon entropy of the cluster assignment. We then regressed the diversity score against age to detect the systematic increase/decrease of clustering diversity with respect to age. FDR was used to correct for multiple comparisons. A tissue or a tissue-cell type was selected if the slope was consistent (having the same sign) in all 6 clustering configurations and at least 2 out of 6 clustering configurations had FDR<0.3. For each selected tissue or tissue-cell type, a separate UMAP was computed using cells from all mice for visualization using Leiden clustering with resolution parameter 0.7.

## Code availability

All code used for analysis is available on GitHub (https://github.com/czbiohub/tabula-muris-senis)

## Interactive Data Browsers

tabula-muris-senis.ds.czbiohub.org

https://tabula-maris-senis.cells.ucsc.edu

**Supplementary Information** is available in the online version of the paper.

## Acknowledgements

We thank Sony Biotechnology for making an SH800S instrument available for this project. Some cell sorting/flow cytometry analysis for this project was done on a Sony SH800S instrument in the Stanford Shared FACS Facility. Some fluorescence activated cell sorting (FACS) was done with instruments in the VA Flow Cytometry Core, which is supported by the US Department of Veterans Affairs (VA), Palo Alto Veterans Institute for Research (PAVIR), and the National Institutes of Health (NIH). We would like to thank Bruno Tojo for helping with the artwork. We thank Chenling Antelope and Joshua Batson for helpful discussions.

## Author Contributions

### Overall Coordination

Angela Oliveira Pisco^1^, Aaron McGeever^1^, Nicholas Schaum^2^, Jim Karkanias^1^, Norma F. Neff^1^, Spyros Darmanis^1*^, Tony Wyss-Coray^3-5*^, and Stephen R. Quake^1,6*^

* Correspondence to

### Organ collection and processing

Jane Antony^2^, Ankit S. Baghel^2^, Isaac Bakerman^2,7,8^, Ishita Bansal^3^, Daniela Berdnik^5^, Biter Bilen^3,4^, Douglas Brownfield^9^, Corey Cain^10^, Michelle B. Chen^4^, Stephanie D. Conley^2^, Spyros Darmanis^1^, Aaron Demers^2^, Kubilay Demir^2,11^, Antoine de Morree^3,4^, Tessa Divita^1^, Haley du Bois^5^, Laughing Bear Torrez Dulgeroff^2^, Hamid Ebadi^1^, F. Hernán Espinoza^9^, Matt Fish^2,11,12^, Qiang Gan^3,4^, Benson M. George^2^, Astrid Gillich^9^, Foad Green^1^, Geraldine Genetiano^1^, Xueying Gu^12^, Gunsagar S. Gulati^2^, Yan Hang^12^, Shayan Hosseinzadeh^1^, Albin Huang^4^, Tal Iram^4^, Taichi Isobe^1^, Feather Ives^2^, Robert Jones^3^, Kevin S. Kao^2^, Guruswamy Karnam^13^, Aaron M. Kershner^2^, Nathalie Khoury^3^, Bernhard M. Kiss^2,14^, William Kong^2^, Maya E. Kumar^15,16^, Jonathan Lam^12^, Davis P. Lee^6^, Song E. Lee^4^, Olivia Leventhal^5^, Guang Li^17^, Qingyun Li^18^, Ling Liu^3,4^, Annie Lo^1^, Wan-Jin Lu^1,9^, Maria F. Lugo-Fagundo^5^, Anoop Manjunath^1^, Andrew P. May^1^, Ashley Maynard^1^, Marina McKay^1^, M. Windy McNerney^32,33^,Ross J. Metzger^19,20^, Marco Mignardi^1^, Dullei Min^21^, Ahmad N. Nabhan^9^, Norma F. Neff^1^, Katharine M. Ng^3^, Joseph Noh^2^, Rasika Patkar^13^, Weng Chuan Peng^12^, Lolita Penland^1^, Robert Puccinelli^1^, Eric J. Rulifson^12^, Nicholas Schaum^2^, Michael Seamus Haney^3^, Shaheen S. Sikandar^2^, Rahul Sinha^2,22-24^, Rene V. Sit^1^, Daniel Staehli^3^, Krzysztof Szade^2,25^, Weilun Tan^1^, Cristina Tato^1^, Krissie Tellez^12^, Kyle J. Travaglini^9^, Carolina Tropini^26^, Lucas Waldburger^1^, Linda J. van Weele^2^, Michael N. Wosczyna^3,4^, Jinyi Xiang^2^, Soso Xue^3^, Andrew C. Yang^3^, Lakshmi P. Yerra^5^, Justin Youngyunpipatkul^1^, Fabio Zanini^3^, Macy E. Zardeneta^6^, Fan Zhang^19^, Hui Zhang^5^, Lu Zhou^18^

### Library preparation and sequencing

Spyros Darmanis^1^, Shayan Hosseinzadeh^1^, Ashley Maynard^1^, Norma F. Neff^1^, Lolita Penland^1^, Rene V. Sit^1^, Michelle Tan^1^, Weilun Tan^1^, Alexander Zee^1^

### Computational Data Analysis

Oliver Hahn^3^, Lincoln Harris^1^, Andreas Keller^3,36^, Benoit Lehallier^3^, Aaron McGeever^1^, Angela Oliveira Pisco^1^, Róbert Pálovics^3^, Weilun Tan^1^, Martin Jinye Zhang^30,37^

### Cell Type Annotation

Nicole Almanzar^21^, Jane Antony^2^, Biter Bilen^3,4^, Weng Chuan Peng^12^, Spyros Darmanis^1^, Antoine de Morree^3,4^, Oliver Hahn^3^, Yan Hang^12^, Mu He^31^, Shayan Hosseinzadeh^1^, Tal Iram^4^, Taichi Isobe^1^, Aaron M. Kershner^1^, Jonathan Lam^12^, Guang Li^17^, Qingyun Li^18^, Ling Liu^3,4^, Wan-Jin Lu^1,9^, Ashley Maynard^1^, Dullei Min^21^, Ahmad N. Nabhan^9^, Patricia K. Nguyen^2,7,8,17^, Angela Oliveira Pisco^1^, Zhen Qi^2^, Nicholas Schaum^2^, Joe M. Segal^13^, Shaheen S. Sikandar^2^, Rahul Sinha^1,22-24^, Rene Sit^1^, Michelle Tan^1^, Weilun Tan^1^, Kyle J. Travaglini^9^, Margaret Tsui^13^, Bruce M. Wang^13^, Linda J. van Weele^2^, Michael N. Wosczyna^3,4^, Jinyi Xiang^2^, Alexander Zee^1^, Lu Zhou^18^

### Liver staining and data analysis

Rafael Gòmez-Sjöberg^1^, Angela Oliveira Pisco^1^, Joe M. Segal^13^, Margaret Tsui^13^, Kevin A Yamauchi^1^

### Microbiome analysis

Bryan Merrill^26^, Aaron McGeever^1^, Katharine M. Ng^3^, Angela Oliveira Pisco^1^, Carolina Tropini^26^, Brian Yu^1^, Chunyu Zhao^1^, Katherine Pollard^34^, Justin Sonnenburg^26^, Kerwyn Casey Huang^2,3^

### Writing Group

Spyros Darmanis^1^, Angela Oliveira Pisco^1^, Stephen R. Quake^1,6^, Tony Wyss-Coray^3-5^

### Principal Investigators

Ben A. Barres^18^, Philip A. Beachy^2,9,11,12^, Charles K. F. Chan^28^, Michael F. Clarke^2^, Spyros Darmanis^1^, Kerwyn Casey Huang^2,3,26^, Jim Karkanias^1^, Seung K. Kim^12,29^, Mark A. Krasnow^9,11^, Maya E. Kumar^15,16^, Christin S. Kuo^9,11,21^, Ross J. Metzger^19,20^, Norma F. Neff^2^, Roel Nusse^9,11,12^, Patricia K. Nguyen^2,7,8,17^, Thomas A. Rando^3-5^, Justin Sonnenburg^26^, Bruce M. Wang^13^, Kenneth Weinberg^21^, Irving L. Weissman^2,22-24^, Sean M. Wu^2,7,17^, James Zou^1,30,35^, Stephen R. Quake^2,6^, Tony Wyss-Coray^3-5^

^1^ Chan Zuckerberg Biohub, San Francisco, California, USA

^2^ Institute for Stem Cell Biology and Regenerative Medicine, Stanford University School of Medicine, Stanford, California, USA

^3^ Department of Neurology and Neurological Sciences, Stanford University School of Medicine, Stanford, California, USA

^4^ Paul F. Glenn Center for the Biology of Aging, Stanford University School of Medicine, Stanford, California, USA

^5^ Center for Tissue Regeneration, Repair, and Restoration, V.A. Palo Alto Healthcare System, Palo Alto, California, USA

^6^ Department of Bioengineering, Stanford University, Stanford, California, USA

^7^ Stanford Cardiovascular Institute, Stanford University School of Medicine, Stanford, California, USA

^8^ Department of Medicine, Division of Cardiology, Stanford University School of Medicine, Stanford, California, USA

^9^ Department of Biochemistry, Stanford University School of Medicine, Stanford, California, USA

^10^ Flow Cytometry Core, V.A. Palo Alto Healthcare System, Palo Alto, California, USA

^11^ Howard Hughes Medical Institute, USA

^12^ Department of Developmental Biology, Stanford University School of Medicine, Stanford, California, USA

^13^ Department of Medicine and Liver Center, University of California San Francisco, San Francisco, California, USA

^14^ Department of Urology, Stanford University School of Medicine, Stanford, California, USA

^15^ Sean N. Parker Center for Asthma and Allergy Research, Stanford University School of Medicine, Stanford, California, USA

^16^ Department of Medicine, Division of Pulmonary and Critical Care, Stanford University School of Medicine, Stanford, California, USA

^17^ Department of Medicine, Division of Cardiovascular Medicine, Stanford University, Stanford, California, USA

^18^ Department of Neurobiology, Stanford University School of Medicine, Stanford, CA USA

^19^ Vera Moulton Wall Center for Pulmonary and Vascular Disease, Stanford University School of Medicine, Stanford, California, USA

^20^ Department of Pediatrics, Division of Cardiology, Stanford University School of Medicine, Stanford, California, USA

^21^ Department of Pediatrics, Stanford University school of Medicine, Stanford, California, USA

^22^ Department of Pathology, Stanford University School of Medicine, Stanford, California, USA

^23^ Ludwig Center for Cancer Stem Cell Research and Medicine, Stanford University School of Medicine, Stanford, California, USA

^24^ Stanford Cancer Institute, Stanford University School of Medicine, Stanford, California, USA

^25^ Department of Medical Biotechnology, Faculty of Biophysics, Biochemistry and Biotechnology, Jagiellonian University, Poland

^26^ Department of Microbiology & Immunology, Stanford University School of Medicine, Stanford, California, USA

^27^ Department of Biochemistry and Biophysics, University of California San Francisco, San Francisco, California USA

^28^ Department of Surgery, Division of Plastic and Reconstructive Surgery, Stanford University, Stanford, California USA

^29^ Department of Medicine and Stanford Diabetes Research Center, Stanford University, Stanford, California USA

^30^ Department of Electrical Engineering, Stanford University, Palo Alto, 94304 USA

^31^ Department of Physiology, University of California, San Francisco, CA 94158

^32^ Mental Illness Research Education and Clinical Center, V.A. Palo Alto Healthcare System, Palo Alto, California, USA

^33^ Department of Psychiatry, Stanford University School of Medicine, Stanford, California, USA

^34^ Department of Epidemiology and Biostatistics, University of California, San Francisco, CA 94158

^35^ Department of Biomedical Data Science, Stanford University, Palo Alto, 94304 USA

^36^ Clinical Bioinformatics, Saarland University, Saarbrücken, Germany

^37^ Department of Epidemiology, Harvard T.H. Chan School of Public Health, Boston, Massachusetts, USA

